# Peptidome and Metabolome profiling of the fermented milk products revealed accumulation of bioactive compounds during two weeks of cold storage

**DOI:** 10.1101/2025.07.11.664382

**Authors:** Elena Kovalenko, Maxim Sheludchenko, Olesya Shoshina, Vera Odintsova, Stanislav Koshechkin, Olesya Volokh

**Affiliations:** Health&Nutrition LLC, Moscow region, Krasnogorsk city district, Russia; Nobias Technologies LLC, Moscow, Russia

**Author notes:** Corresponding author: Maxim Sheludchenko, Moscow region, Krasnogorsk city district, Riga-Land Business Centre, building 4, H&N office, Russia. All work was funded by Health&Nutrition LLC.

**Keywords:** peptidome, fermented milk, kefir, metabolome, lipidome

## Abstract

This study assessed the metabolome and peptidome profiles of various fermented milk products produced using different starter cultures. Milk fermentation involves a set of macromolecular decomposition reactions performed by microorganisms under controlled conditions. Biotransformation of fats, proteins, and carbohydrates shapes the metabolome profile of each fermented product. Many of these microbe-generated molecules exhibit biological activities that can affect human health. The resulting profile is unique for every fermented product and depends on technology, raw materials, and starter culture composition. We used four non-targeted metabolomic methods to assess semi-quantified concentrations of four types of molecules in the final products: peptides; amino acids; long-, medium-, and short-chain fatty acids; mono- and disaccharides and their derivatives. Ultra-performance liquid chromatography-mass spectrometry (UPLC-MS/MS) was performed on the peptidome. For all other fractions, we used gas chromatography–mass spectrometry (GC-MS), with a method adapted to specific metabolite conditions. Metabolome and peptidome of four groups of 15 dairy cow milk products including yogurt (Y), fermented milk (FM), kefir made with commercial cultures (K) and kefir made with grains (KG) was performed on days 7 and 14 of shelf life at 4°C, and milk (M) was used as a control. Peptides, amino acids, fatty acids, mono- and disaccharides, and their low molecular weight derivatives were evaluated. In total, 348 peptides, 37 amino acids, 25 fatty acids, and 23 mono- and disaccharides were identified in the products. Among them, 41 functional peptides, branch-chained amino acids (BCAA), orotic acid (vitamin B13), d-phenyllactic acid, 5-phenylvaleric acid and myo-inositol (vitamin B8) accumulated in fermented products during storage.

## 1. Introduction

Fermentation is an ancient way for preserving food and improving its shelf life ^1,2^. In recent decades, fermented milk products have garnered considerable research attention owing to their numerous health benefits, such as improved digestion and bioavailability of milk constituents, inhibition of harmful gastrointestinal bacteria, alleviation of lactose intolerance, and effects of fermented milk products and gut microbiome on brain activity; therefore, they are considered functional foods ^3–6^. Metabolites reflect the quality of milk and dairy products in terms of their nutritional value and safety ^7^. Notably, the identified differentially abundant metabolites are believed to play a crucial role in shaping the distinctive gustatory and olfactory characteristics of fermented milk products ^8^. Metabolism of macronutrients is essential for the production of fermented milk products such as yogurt, cheese, and kefir. The specific types and quantities of metabolized compounds can be affected by the microorganisms present in the starter culture. Factors such as raw milk, fat percentage, season, temperature, and geographical origin as well as contamination or adulteration during processing and transport may also alter metabolite profiles produced by microorganisms ^9–12^.

In the case of proteins, fermentation involves microbial hydrolysis of peptide bonds, which results in the release of oligopeptides and free amino acids. The initial amount of free amino acids and peptides in milk is limited and only sustains the starter culture for up to five generations ^13^. Therefore, lactic acid bacteria (LAB) and yeasts employ a complex proteolytic system to degrade milk proteins. It has been observed that the proteolytic activity in milk is associated with the formation of polypeptides and oligopeptides through the action of bacterial peptidases that break down larger molecules, resulting in the production of smaller peptides and free amino acids ^8^. Studies have also shown that bacterial proteases and peptidases remain active during fermentation and storage of fermented milk and these enzymes exhibit specificity for specific cleavage sites or sequences during proteolysis ^14^.

Fats in fermented food are broken down by lipolysis. The medium-length fraction of fatty acids appears to be utilized by microorganisms during their metabolic processes, thereby preventing their excessive accumulation ^15^. The fraction of short fatty acids is the most prominent component of fermented products ^16^. This could be a consequence of lipolysis and glycolysis. Through these mechanisms, the fat fraction undergoes transformations, leading to varied compositions of the different fermented products. LAB are thought to have weak lipolytic activity compared to yeasts or other bacterial taxa such as *Pseudomonas*, *Flavobacterium,* and *Staphylococcus* ^17,18^. However, in products such as cheese with extended ripening times, the lipolytic activity of LAB contributes to flavor development and serves as a substrate for further reactions, leading to the formation of end-products ^19^. Therefore, the evaluation of the degree of lipolysis helps might aid in the design of to determine the choice of strains used as starter cultures.

Carbohydrates are other potential substrates in milk that can be readily used by microorganisms. LAB secrete various extracellular and capsular polysaccharides. Complex carbohydrates play a crucial role in determining the unique textural properties of various fermented milk products, influencing their viscosity, creaminess, and mouthfeel. Dairy oligosaccharides are composed of 3–20 monosaccharides ^20^. Typically only 30% of lactose is converted into glucose and galactose by β-galactosidase during milk fermentation ^21^. Glucose is metabolized by bacteria into lactic acid via glycolysis ^22^.

Knowledge of the biochemical composition of dairy products with various combinations of starter cultures can provide clues about their nutritional value and opens new opportunities for dietary applications, and some metabolites can be used to ensure quality and safety. Metabolomics has emerged as a powerful tool for understanding the intricate metabolic processes that occur during fermentation, to further unravel the complex biochemical transformations underlying these benefits for food products with improved functionality and health benefits ^23^.

In the current study, we focused on two aims: to evaluate the potentially beneficial major organic compounds of fermented dairy products in contrast to raw milk and to assess the decomposition dynamics of compounds with high-molecular weight during cold storage. The obtained outputs of peptidome, free fatty acids, lipidome, and glycome analyses revealed the accumulation of new functional peptides, fatty acids, BCAAs, and vitamins B8 and B13 over a period of two weeks.

## 2. Material and Methods

### 2.1.1. Samples description, storage characteristics and aliquots taken for the analysis

The current study was carried out on a range of fermented milk drinks: kefir made with wild kefir grains (KG) with 1.2% and 2.5% fat (n=2); kefir drink (K) made with commercial cultures of *Lactobacillus acidophilus, Lactococcus lactis, Lactococcus mesenteroides, Bifidobacterium animalis* and *Debaryomyces hansenii* with 0.1% and 1% fat (n=2), fermented milk (FM) with 2.5% fat (n=1); probiotic yogurt (Y) with 1.7% fat (n=1); and normalized milk (M) by 1.2%, 2.5%, 3.2% (n=3) fat taken as a reference without fermentation. All products were supplied by the dairy company Health&Nutrition LLC. The concentrations of LAB in the starter cultures were at least 1x10^10^ CFU/g, *B. animalis* at least 1x10^8^ CFU/g and yeast at least 1x10^4^ CFU/g. K contained a single yeast strain, KG contained 10 yeast strains in the starter cultures and FM and Y did not contain any yeast in the starter cultures. Each product with a range of various fat contents (except milk) was sampled at 7 and 14 days of storage at +2 ..+4°C. To study the decomposition dynamics of the substances, we used these two time points for the shelf life (D+7 vs. D+14). In total, 15 dairy products were analysed (Table 1).

**Table 1.**
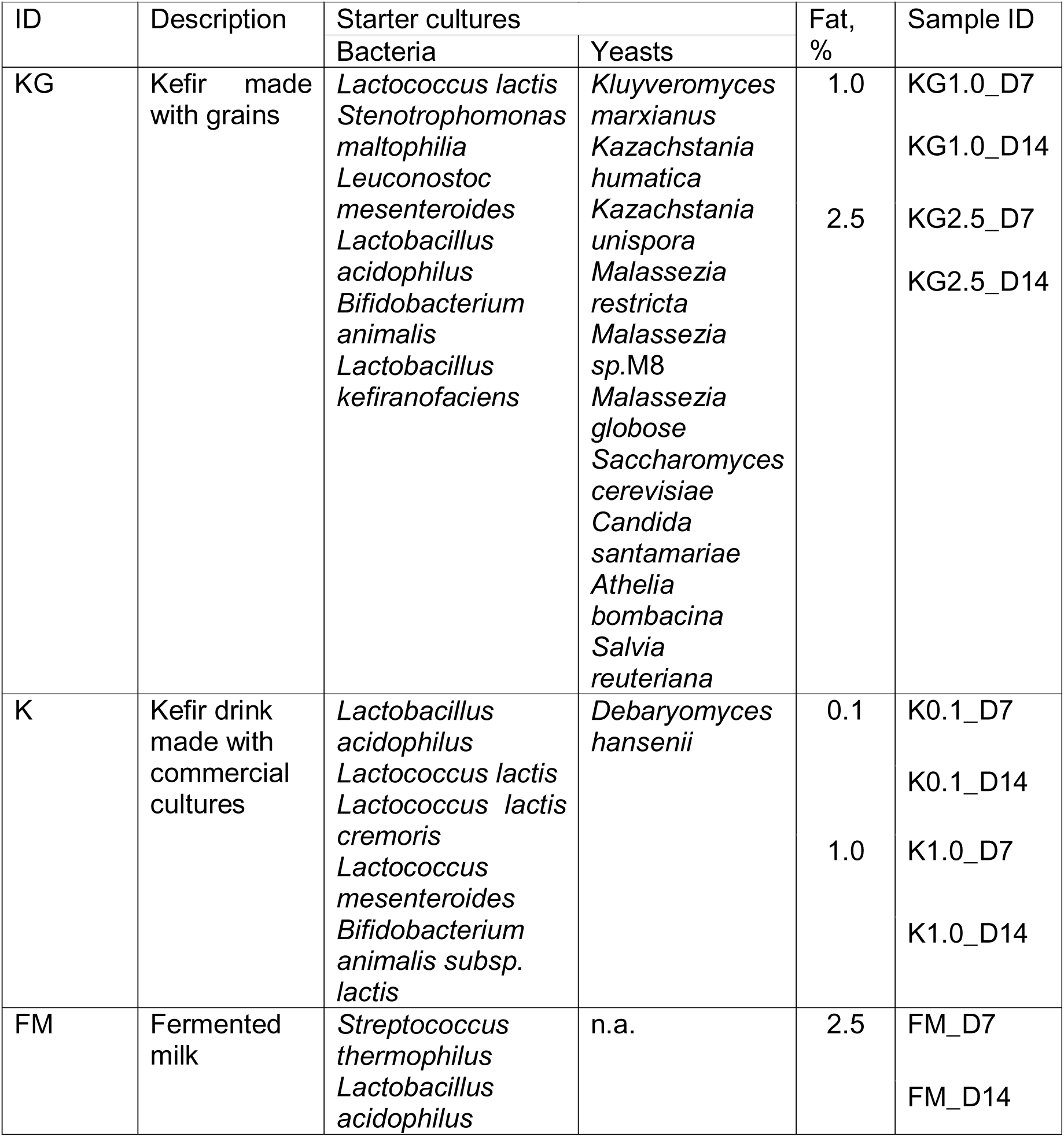
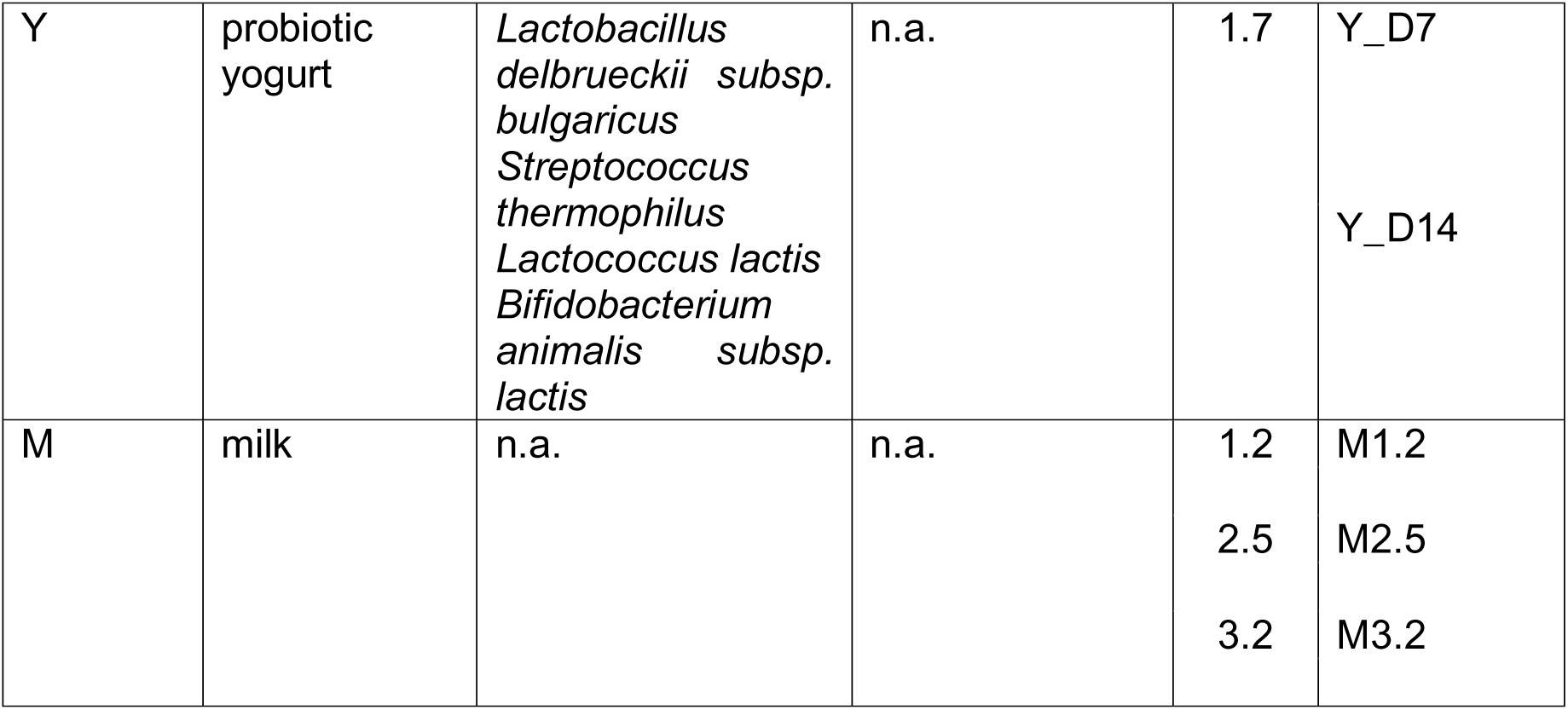
Microbial composition and fat content of the products used in this study. To study the decomposition dynamics of these substances, two samples were collected on days 7 (D7) and 14 (D14). Only microbial species with > 1% abundance are shown for the KG samples.

For each dairy product, 250 mL of sample in five aliquots of 50 mL were used for further analysis. The samples were stored at -35 °C until brought into the laboratory.

Metabolites analyses were used to evaluate peptidome and four varieties of untargeted metabolomes to evaluate different fractions of organic compounds (fatty acids, short fatty acids, amino acids, and sugars) (Fig. 1). In addition to metabolite analysis, biochemical methods were used to determine total protein in the sample and total free peptides as part of the sample preparation, as well as total fat (triglycerides). The description of the methods used in this study is presented below.

**Figure 1.**
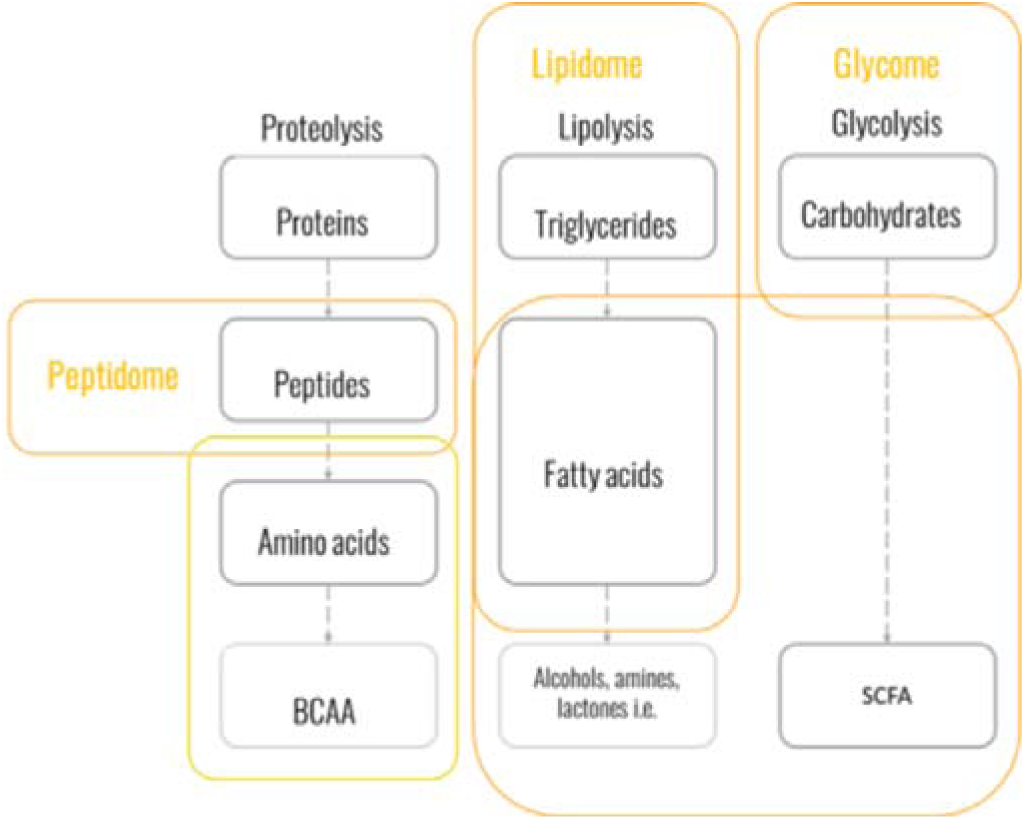
Schematic representation methods (peptidome, free amino acids analysis, lipidome and glycome) applied in this study for peptidome and metalobome profiling.

### 2.1.2. 16s rRNA and ITS amplicon sequencing analysis of KG samples

Samples of kefir grains and white mass of KG were received in triplicate (20 mL each) in a frozen format and preserved at -20°C until processed. DNA was extracted using an internal protocol with the aid of a MagnaPure automatic extractor (Roche Life Sciences). Two sets of primers were used, with the first pair capturing region for the 16s rRNA ^24^ of the bacterial taxa, and the second pair capturing region for ITS ^25^ of the fungi. Artificial mock community, composed of 25 *Lactobacillus* and 75 *Bifidobacterium* strains was included as a positive control. Samples were sequenced at Biopolis S.L. CIFB (Valencia, Spain) facility with MiSeq Platform (Illumina, USA) using a 300PE combination.

For bioinformatic analysis, pair-end raw sequences were merged in order to obtain the complete sequence with ’pear v0.9.6’. The amplification primers were trimmed by ’cutadapt v 1.8.1’ with default parameters. Sequences less than 200 nt were not further considered for the analysis and a quality filter was applied to delete sequences of poor quality. Bases in extreme positions that did not have Q20 (99%) of well incorporated bases in the sequencing step or more pared score were removed, sequences which quality mean did not surpass the Q20 threshold, as a mean quality of the whole sequence, were also deleted. The resulting sequences were inspected for PCR chimera constructs that may occur during different experimental processes and chimeras were removed. In order to reduce the complexity of the annotation, sequences with a 97% of similarity were clustered into a single cluster using program ‘cd-hit’, so only the representative sequences need to be annotated, and results were applied to the cluster of sequences represented by the analysis. Each of the clean clustered sequences were compared against the built rRNA database using a BLAST local alignment approach based on publicly available source https://ftp.ncbi.nlm.nih.gov/blast/db/, in order to associate each of the cluster to one taxonomical group from the database.

### 2.2. Peptidome analysis

#### 2.2.1. Sample preparation for peptidome by free peptide extraction

Three biological replicates of 10 mL from 50 mL thawed aliquot were lyophilized in 50 mL tubes. After, 20 µg lyophilized dried mass from each sample were resuspended into 1 mL of 3% acetonitrile (ACN) and 0.1% trifluoroacetic acid (TFA) (Sigma-Aldrich, cat#1.08178) in milli-Q water (Millipore ZLXE0030RU) and shaken at 2000 rpm (ELMI S-3M. A10) for 30 min. After centrifugation at 1100 rpm for 30 min, the supernatant was collected and filtered through a 0.22 µm 25 mm FLL/mIS CA ST syringe filter (GVS, Italy). The obtained extracts were further purified on using a C18 column Oasis (Waters, US). The eluates were dried under vacuum and diluted into 30 µL milli-Q water containing 3% ACN and 0.1% TFA. The peptide concentration was determined using the BCA Assay Kit (Sigma-Aldrich, cat #71285-M) for all techniques.

#### 2.2.2. Ultra-performance Liquid Chromatography-mass spectrometry analysis (LC/MS)

Free peptides extracts were analysed separately using a nano-ESI Orbitrap Q Exactive HF-X mass spectrometer (Thermo Fisher Scientific, Waltham, MA, USA) coupled to a nano-flow high-pressure chromatograph (UPLC Ultimate 3000) with a reverse-phase C18 column 100 µm x 300 mm (Thermo Fisher Scientific, USA). For sample analysis, a gradient of 0-40% buffer B (buffer A: 0.1% formic acid in DI water (18Mohm) and buffer B: 80% acetonitrile and 0.1% formic acid in DI water) was used with a flow rate of 0.25 µl/min. Emitter needle voltage was set to +2.2 kV, RF voltage at S-lens was 65, capillary temperature was 250°C. The MS and MS/MS spectra were acquired at resolutions of 60,000 and 30,000 respectively. Charge accumulation cut off levels were 2e6 for MS and 2e5 for MS/MS mode with maximum ion accumulation times of 45 msec and 50 msec. The protocol for the assessment of relative peptide concentrations can be found in Supplementary Materials 2.

### 2.3. Lipidome, glycome and free amino acids analysis

#### 2.3.1. Total triglycerides analysis

Sample preparation was performed using two biological replicates. Seven grams of each sample were thawed with 7 mL of concentrated HCl, mixed and kept in an oven at 90°C for 1 h. After cooling to room temperature, samples were mixed with 2 mL methanol and 20 mL hexane, shaken vigorously on vortex and the hexane layer was collected into new tube. The remaining water fraction of the same sample was mixed with 2 mL methanol, 20 mL diethyl ether was added, shaken vigorously on vortex and obtained extract was combined with hexane extract. Remaining watery fraction was mixed with 500 mg NaCl, extracted with 20 mL hexane and combined with other extracts. The combined final extracts were dried in an oven at 90°C to obtain a constant weight. The percentage of fat in the original sample was calculated based on difference between the initial weight and weight of the dried sample.

#### 2.3.2. Free fatty acids analysis

Thawed samples (20 g) from two biological replicates were mixed with 10 g of NaCl on an orbital shaker. Once salt was completely dissolved, 25 mL ACN and 1 mL of 10% HCl were added to get pH 1-2. Samples were incubated at room temperature with 450 rpm for 10 min, followed by centrifugation at 3500 rpm for 10 min. The supernatant was collected, mixed with 20 mL of water, neutralized with 5M KOH to get pH 5-6 and obtained mix was evaporated to 20 mL. Next, 5M KOH was added to get pH 10 and added 25 mL of hexane. The mixture was incubated at room temperature on a shaker with 450 rpm during 10 min and then centrifuged at 3500 rpm for 10 min. The bottom aqueous layer was collected, acidified with 10% HCl to get pH 2-2.5, mixed with 25 mL dichloromethane and incubated at room temperature with 450 rpm for 10 min, followed by centrifugation at 3500 rpm for 10 min. The bottom layer was collected and ACN layer was collected and evaporated to a minimal volume, transferred into glass vial and dried under nitrogen gas (Microvap, Oranomation, MA, US) at 60°C. The residue was derivatized with 70 µL of N-Methyl-N-(trimethylsilyl)-trifluoroacetamide (MSTFA) (Sigma-Aldrich, cat # 69479), incubated at 70°C for 30 min, mixed with 400 µL of ethyl acetate and analysed by GC-MS.

#### 2.3.3. Free carbohydrates (mono- and disaccharides) analysis

Thawed samples in two biological replicates (5 g fermented milk or 1 g milk) were extracted with ACN and acidified with 10% HCl to get a pH of 1-2 by mixing on an orbital shaker at room temperature for 10 min. Protein fractions were separated by centrifugation at 3500 rpm for 10 min. Bottom aqueous layer of 1 mL was transferred into glass vial and neutralized with 5M KOH to get pH 5-6, evaporated to minimal volume at heater block and completely dried with addition of isopropanol under nitrogen flow at 60°C. After, pellet was derivatized with 200 µL MSTFA and incubated under a closed cap at 70°C for 20 h. The derivatized carbohydrates were mixed with 400 µL ethyl acetate, vortexed, transferred into a 1.5 mL tube and centrifuged at 13500 rpm for 10 min at room temperature. Supernatant was then placed into a glass vial for GC-MS analysis.

#### 2.3.4. Free amino acids analysis

Sample preparation was performed independently in two biological replicates. Ten grams of NaCl were added to 20 g of the thawed sample, and the mixture was stirred on a shaker until the salt was completely dissolved. Next, 25 mL of ACN was added and acidified with 1.0 mL of 10% HCl to get pH 1-2. Mix was incubated at room temperature with stirring at 450 rpm for 10 min and centrifuged at 3500 rpm for 10 min. Obtained upper layer with ACN was mixed with 15 mL of water and 5M KOH solution to get pH 5-6, then evaporated to 20 mL. A solution of 5M KOH was added to get pH 10, and 20 mL of hexane was added and the mixture was incubated at room temperature with stirring at 450 rpm for 10 min, followed by centrifugation at 3500 rpm for 10 min.

Bottom aqueous layer was collected and acidified with 10% HCl to pH 2-2.5. Next, 25 mL of dichloromethane was added and incubated at room temperature at 450 rpm for 10 min followed by centrifugation at 3500 rpm for 10 min. The collected upper aqueous layer was evaporated to a minimum volume in a glass vial and five times volume of isopropanol was added and dried under nitrogen flow at 60°C. The dried sample was mixed with 100 μL of MSTFA and the reaction was carried out under a closed cap at 70°C for 4 h and then mixed with 400 µL of ethyl acetate. Finally, the mixture was centrifuged at 13500 rpm for 10 min at room temperature and supernatant was transferred to a 1.5 mL vial for GC/MS analysis.

#### 2.3.5. Gas chromatography with mass-spectrometry (GC-MS)

Compounds profiles for the samples were acquired with use of Agilent 8890 GC System (Agilent, Singapore) with mass-spectrometry detector Agilent 5977B GC/MSD equipped with Agilent 7693 autosampler. Analytes were separated with HP-5MS column (30 m x 0.25 mm x 0.25 µm) (19091S-433UI-KEY, Agilent, USA) under helium at pressure of 95 kPa and gas flow 1.3 µL/min Volume of injection was 1 µL with conditions specified for class of compounds.

For free fatty acids, following conditions were used: injection at 250°C without flow division. Initial oven temperature was 100°C for 5 min with a final temperature of 280°C, a ramping rate 4°C/min, dwell time was 12 min with overall analysis 62.0 min. MS detector temperature was 280°C with MS in full scanning mode in the range of 40-700 m/z with a frequency of 12.8 scans /sec.

For free carbohydrates, the following conditions were used: injection at 270°C, with a flow separation of 5:1. Initial oven temperature was 100°C for 1 min with final temperature 300°C with ramping rate 25°C/min, dwell time was 5 min at 250°C with overall time of analysis 14.0 min. The MS detector temperature was 280°C, with MS in full scanning mode in the range of 40-500 m/z with a frequency of 12.8 scans /sec. For free amino acids, the following conditions were used: injection at 270°C, without gas flow separation. The initial oven temperature was 80°C for 7 min with a final temperature of 100°C; 10 min dwell time at 100°C with ramping rate 3°C/min up to 120°C; 10 min dwell time at 120°C with ramping 3°C/min up to 140°C; 10 min dwell time at 140°C, ramping 3°C/min up to 160°C, 10 min stop-over at 160°C, ramping 3°C/min up to 180°C; 1 min dwell time at 180°C, ramping 4°C/min up to 280°C; and dwell time 4 min at 280°C, with an overall analysis time of 110.3 min. The MS detector temperature was 280°C, with MS in full scanning mode in the range of 40-700 m/z with a frequency of 12.8 scans /sec.

Peaks were analyzed using MassHunter GC/MSD ChemStation software. Untargeted metabolites were identified by comparing the spectra of each peak using the NIST library collection (NIST, Gaithersburg, MD, USA). The linear index difference max tolerance was set to 10, and the minimum matching spectra library search was set to 85% (level 2 identification, as described by the Metabolomics Standards Initiative [MSI]) ^26^.

Each metabolite peak area was normalized to the internal standard of 2-iso-propylmalic acid (Sigma-Aldrich, cat #333115) for carbohydrates and amino acids; Supelco37 standard (Sigma-Aldrich, cat #CRM47885) for free fatty acids followed by generalized log transformation and data scaling by autoscaling (mean-centered and divided by standard deviation of each variable) as described before ^27^. Chromatograms of measurements of metabolites on various days for each type of product are shown in Supplementary Materials 2.

### 2.4. Data analysis

Reported values are the average of two replicates obtained for every sample used for free amino acid, glycome, and lipidome analyses and three replicates were used to obtain average number of relative concentrations reported for peptidome data.

Peptide identification was performed using Proteomicslfq software pipeline 28 (https://github.com/nf-core/proteomicslfq/tree/1.0.0) with the following parameters: input *. raw, --database *.fasta, --add_decoys, --protein_level_fdr_cutoff 0.1, -- max_precursor_charge 7, --enzyme ’unspecific cleavage’, --fixed_mods ’ ’-- search_engines comet. Cow milk protein sequences from the UniProt database ^29^ milk_and_bovine_with_human.fasta were used, the search engine was Comet ^30^: https://uwpr.github.io/Comet/. Two bioactive peptide aggregator databases, BioPepDB ^31^ and MBPDB ^32^, were used to identify peptide function.

Data calculations and visualizations were conducted using R v4.3.3. Heatmaps were generated with heatmap library. The color of the cells indicates the representation of each metabolite after logarithmic transformation and centering with respect to the mean value across all samples, where it was detected using the following formula: clr(x_i) = ln(x_i) - mean(ln(x_1), …. ln(x_n)), where x_i is the measurement of the compound in i-sample; grey cells indicate levels below detection limits for measured compounds. Clustering of metabolites and samples was also performed on normalized values, but centering was performed on all samples, including those where the metabolite was absent, by replacing unknown values with a small number. A cladogram for clustering the samples with important peptides was drawn using the hclust function. Barplots were generated using the ggplot2 and MicrobeR packages. The correlation matrix was obtained with corrplot package, Spearman’s correlation and p-values after Holm’s correction for multiple comparisons were estimated with rcorr.adjust() function from the RcmdrMisc package. The Venn diagram was created with the ggVennDiagram package.

## 3. Results

### 3.1. Peptidome characteristics

Analysis of peptidomes in the 15 products revealed 348 peptides (Table S1). It is evident from the histogram of peptide representation in dairy products that fermented milk products contained significantly more peptides than milk (Fig. 2). Moreover, the peptide composition of 15 products was 75% similar and peptides that were not common to all products were represented at approximately 10 times lower concentrations. When comparing the diversity of peptides in fermented milk products during storage, a slight change was observed, probably due to the limited availability of substrates for peptide formation and the non-optimal conditions for proteolysis (+5°C instead of +37°C).

**Figure 2.**
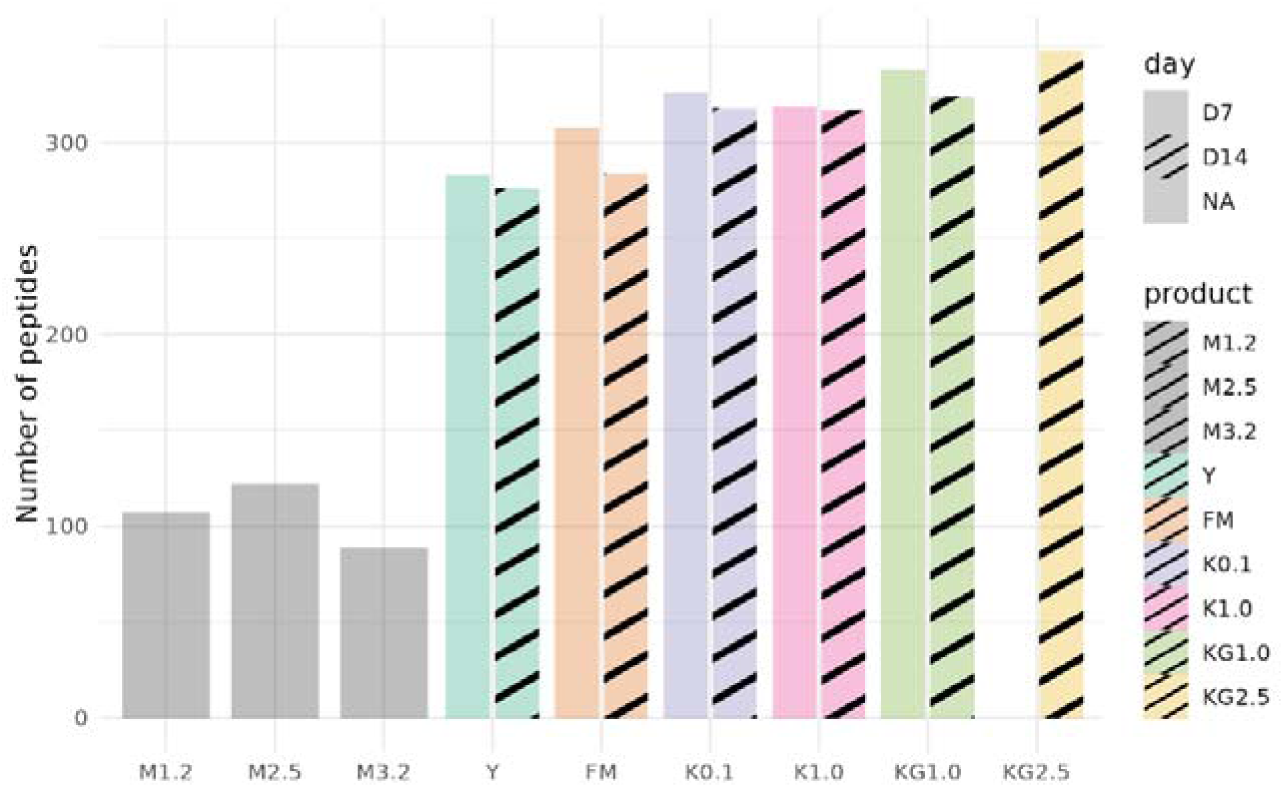
Total number of peptides in different dairy products. KG2.5_D7 sample was excluded from further peptide analysis as it was initially used as a trial sample for the method assessment and was not measured again in batch with other samples. The numbers were calculated based on the unique peptide counts for each product (see Table S6).

The predominant proportion of peptides observed in dairy products (94% on average) originates from the degradation of four proteins: 129 peptides from beta-casein, 72 peptides from kappa-casein, 75 peptides from alpha-S1-casein and 49 peptides from alpha-S2-casein (Fig. 3, Table S1). The remaining peptides were represented by a few peptides and in negligible concentrations relative to the most represented peptides, which might be degradation products of casein proteins and originate from the following proteins of non-casein fractions: whey proteins including major α-lactalbumin (α-LG) and β-lactoglobulin (β-LG); and other proteins such as lactoferrin, lactoperoxidase, osteopontin, perilipin-2, butyrophilin 1/1A and the glycosylation dependent cell adhesion molecule 1 (GLYCAM1).

**Figure 3.**
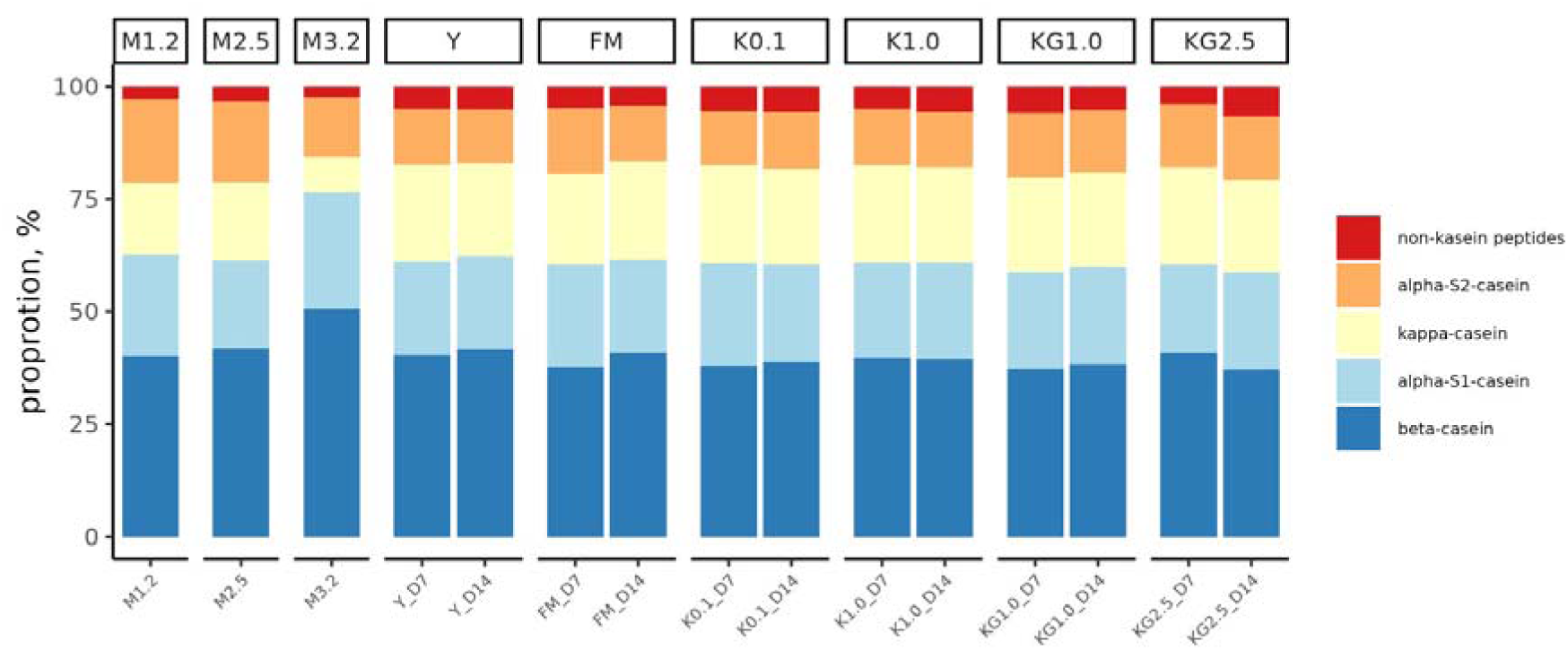
The ratio of the number of peptides derived from various protein fractions.

Based on our study of dairy products, 37.4% of all peptides identified were products of the breakdown of beta-casein, 20.9% of kappa-casein, 21.7% of alpha-S1-casein and 20% of alpha-S2-casein.

When we examined the peptides based on their length and analyzed their accumulation during storage, it was evident that they tended to form clusters according to their size (see Fig. S1-S4). In addition, there was a noticeable increase in the number of smaller peptides over time. This suggests that proteolysis during storage may lead to the generation of shorter peptides by the breakdown of larger peptides or proteins. The formation of peptide clusters and the increase in smaller peptide numbers indicate ongoing proteolytic activity and the dynamic nature of peptide composition during the storage of fermented milk products. It is possible that proteolysis might be localised at various sides of the original protein and the precursors of the target peptide and stop at the point when peptides form an inaccessible steady-state for bacterial protease cleavage. In most cases, these peptides found to be functional ones.

To assess the similarity of peptide composition profiles in fermented milk products, pairwise correlation of samples was performed using peptide composition. A clustering analysis was conducted based on the complete peptide profile (Fig. 4B). Clustering was based on the type of product and resulted in three distinct clusters: M, Y and FM. For the initial time and samples taken after storage, each type of product was more distinct from the other categories than within the same product. This did not apply to kefirs K and KG and their peptidomes did not change significantly during storage. Among the fermented products, FM and Y had the most distinct peptidome profiles. Although Y and FM demonstrated a peptide composition more similar to that of M than K and KG, the level of similarity remained fairly low.

**Figure 4.**
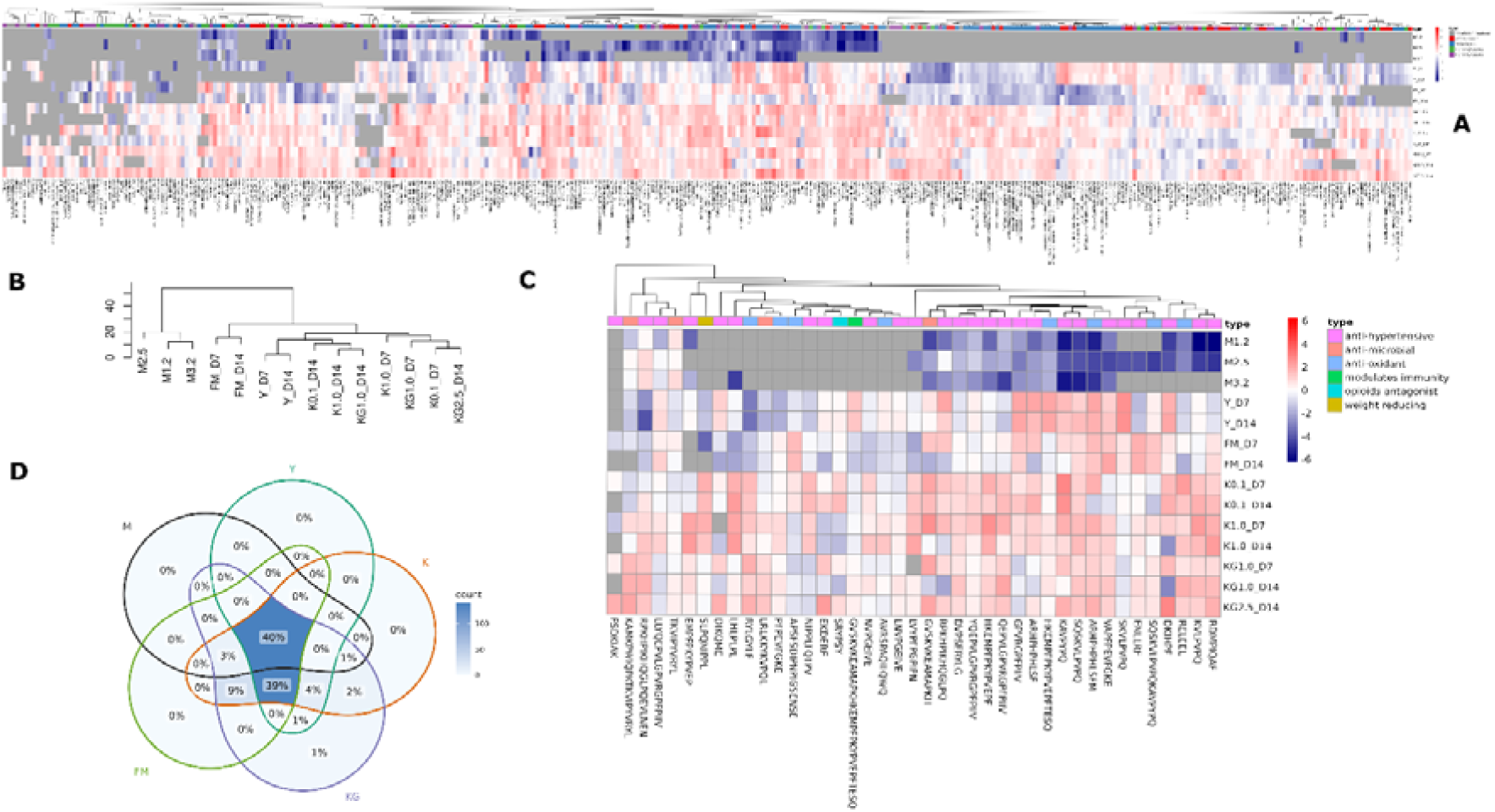
Profiles of 348 peptides for 14 dairy products. KG2.5_D7 was excluded from peptide comparison with other samples as it was analysed in separate batch. (A) Heatmap showing normalized concentration for all peptides identified by untargeted LC/MS, fractions of milk proteins are reflected in the panel and coloured, names of peptides are hidden due to a large number of peptides (Fig. S5). (B) Cluster dendrogram of fermented milk products and milk based on the peptide profiles. (C) Part of the heatmap representing only peptides with known physiological functions in humans. Fermented products are distinct from raw milk. Only peptides identified in fermented products have specific functions such as modulating immunity and stimulating weight reduction. (D) Venn diagram showing the peptides shared between products and milk. Kefir on grains (KG) harbours 1% of distinctive peptides compared to other products. There are three unique peptides that were identified with 1% difference: KEPMIGVN, QEKNMAINPSKE and EPELPLHTL.

This division into clusters may be attributed to the specific characteristics of fermentation and composition of the starter culture. Further interpretation of peptides of the fermented products versus raw milk revealed distinct specific peptide clustering (Fig. 4A) within all analyzed peptides and also identified functional peptides with specific properties that were not found in milk (Fig. 4C). We identified 41 functional peptides from the overall number of peptides reported (see Table S2). Notably, the number of functional peptides in milk was considerably lower than that in any fermented product and products within the same time points were highly similar. Milk peptides appeared to contain low amounts of antihypertensive, antimicrobial, and antioxidant peptides, whereas KG showed the highest enrichment of additional functional peptides responsible for immunity modulation, weight reduction and opioid antagonists. Of note, KG had the highest overall diversity of peptides and 1% of them did not appear in the other products analyzed in this study (Fig. 4D). These included three unique peptides with unknown biological functions: KEPMIGVN, EPELPLHTL and QEKNMAINPSKE.

Most bioactive peptides appear to be unique, but some peptides have been reported previously. We identified the peptide YQEPVLGPVRGPFPIIV in our kefirs (K and KG), which not only exhibited antihypertensive activity but also antithrombotic, immunomodulatory, antioxidative and antimicrobial functions ^33^. This peptide was also found in fermented milk with a single starter culture *Bifidobacterium longum* KACC91563 ^34^ and was produced by a mixed culture of *Streptococcus thermophilus*, *Lb. bulgaricus, Lb. acidophilus, Lb. casei* and *Lb. paracasei* ^35^. Another identified peptide, HKEMPFPKYPVEPF, with antihypertensive functions in all fermented products was previously reported to be a result of proteolysis by mixed cultures of *Lb. acidophilus*, *Lb. delbrueckii* subsp. *bulgaricus* and *Streptococcus thermophilus* ^36^.

We discovered enrichment of the functional peptide DKIHPF with angiotensin inhibitory properties in all the fermented products. Previously, this peptide was reported only in goat milk and appeared to be eight times more active after pepsin digestion ^37^. Peptides GVSKVKEAMAPKHKEMPFPKYPVEPFTESQ and SRYPSY have been previously found to function as agonists and antagonists respectively to opioid receptors in the gut, resulting in mucin production and an impact on peristaltic movement ^38,39^. A single identified SLPQNIPPL peptide has been previously reported to inhibit the activity of dipeptidyl peptidase 4, which is responsible for GLP-1 degradation in the cytosol ^40^.

Interestingly, the occurrence of all oligopeptides analyzed was not found to be related to their origin from a specific protein, nor was it associated with their specific functions. This observation may be attributed to variations in the composition of proteases produced by bacteria and yeast present in the starter culture.

### 3.2. Free amino acids analysis

The assessment of free amino acids was carried out using untargeted metabolomics with evaluation of the obtained peaks relative to library values. A search was conducted for 37 standard compounds that included the majority of biologically significant amino acids including dipeptides (see Table S3). The sensitivity of this metabolomic method was quite low; therefore, only the most abundant analytes were detected. Figure 5 shows that the composition of free amino acids and orotic acid (vitamin B13) differed between milk and fermented products. Moreover, there was a noticeable trend towards an increase in the amount of free amino acids, such as branched chain amino acids (valine, leucine, isoleucine), during storage at D14, despite suboptimal conditions for proteolysis. Accumulation of orotic acid was also observed at D14, whereas its known precursor molecules such as aspartic acid and glutamic acid were depleted. We also detected the accumulation of aromatic amino acids, such as phenylalanine and tyrosine, at D14 with the highest concentrations detected in Y compared with other fermented products. The common amino acids serine, alanine, glycine and tryptophane were not observed in any fermented product even at D7, although they were initially found in milk. This only shows the dynamics of certain amino acids by an active microbial community during cold storage. However, evaluation of absolute values is required for a more complete assessment of the amino acid profile using targeted metabolomic methods in future experiments.

**Figure 5.**
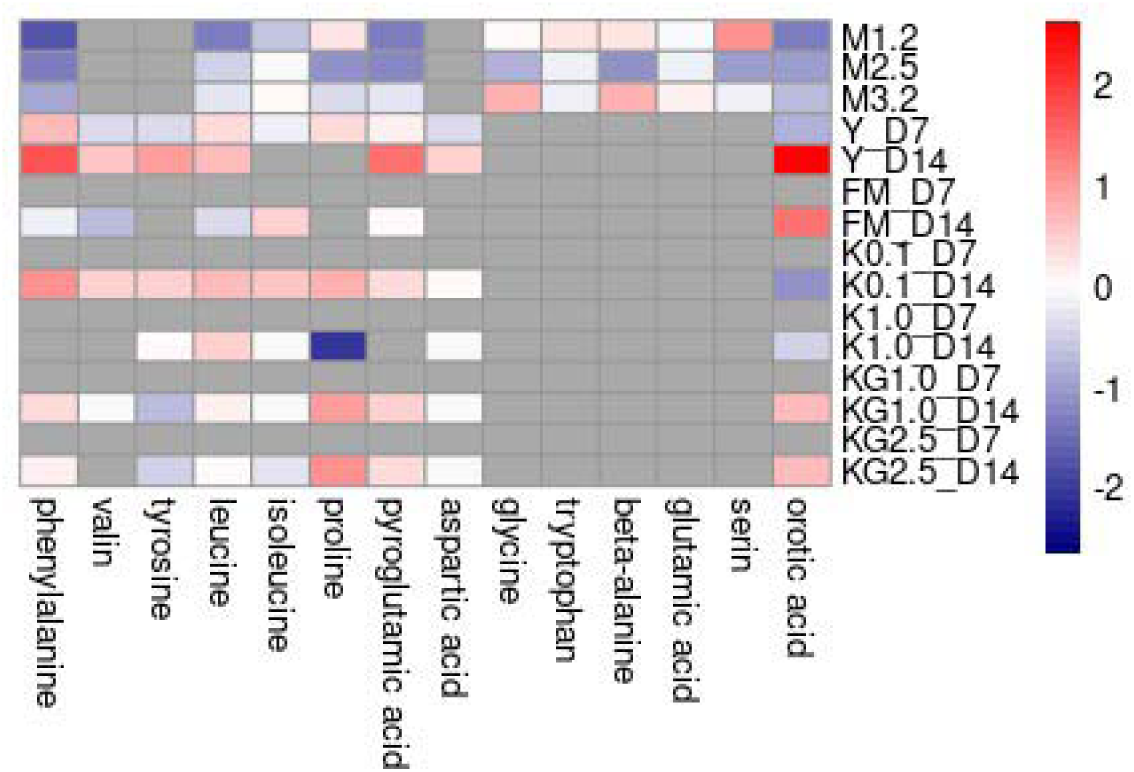
Heatmap of free amino acids and orotic acid concentrations in fermented products and milk

### 3.3. Lipidome analysis

We observed that during the storage of fermented milk products, there was a decrease in the overall number of triglycerides present (Fig. 6). The fat and free fatty acid profiles of all fermented milk products were compared on days 7 and 14 of storage revealing two distinct trends.

**Figure 6.**
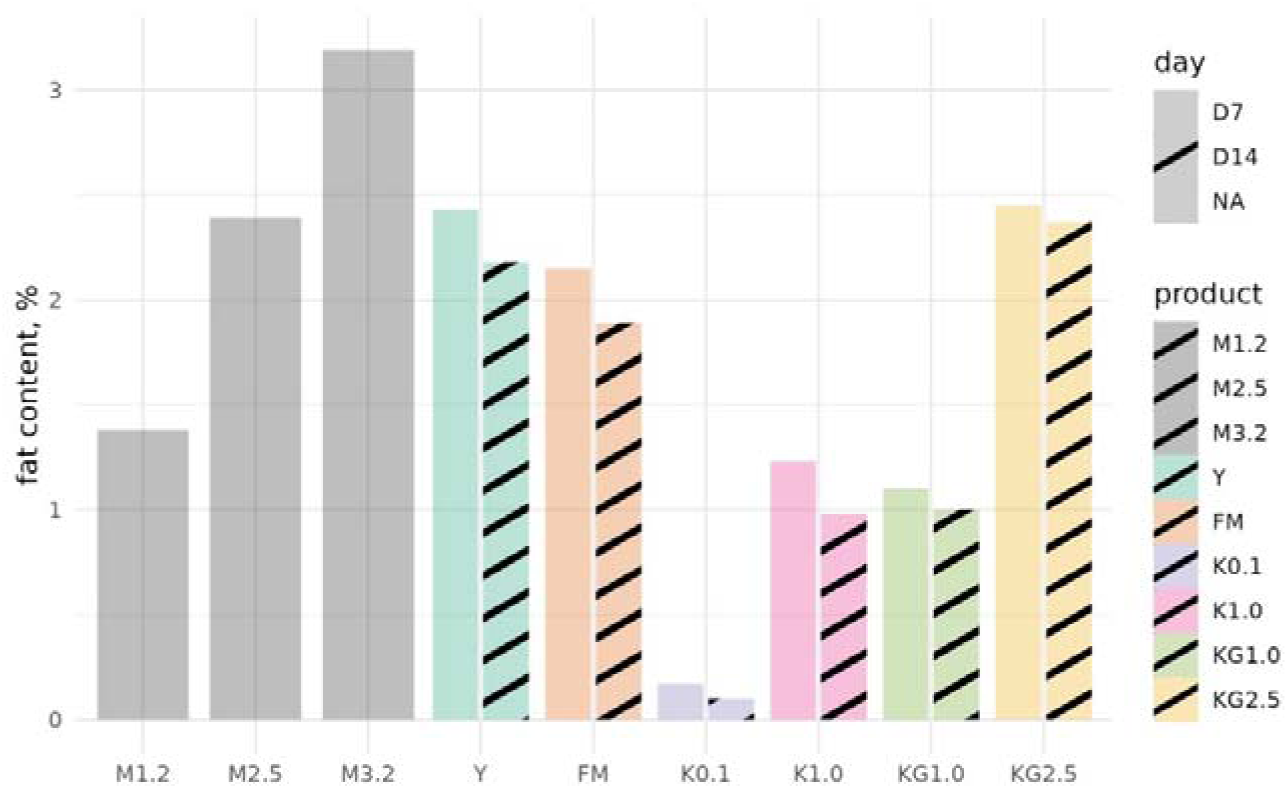
Triglyceride concentration in the milk products based on the fat content (see Table S7).

The table provided (see Table S4) lists the fatty acids detected. . Many of these fatty acids exhibit biological activity and exert positive effects on the human body and microbiome. Among the identified fatty acids, polyunsaturated fatty acids, such as omega-6 and omega-9 have also been identified. In addition to fatty acids, monoglycerides, which are likely products of triglyceride lipolysis, have been identified.

A comparison of the resulting clusters based on the representation of all identified compounds is displayed in the figure below as a heatmap (Fig. 7). Notably, products with a fat content of over 1% exhibit distinctive characteristics of higher medium-chain fatty acid (MCFA) and long-chain fatty acid (LCFA) content.

**Figure 7.**
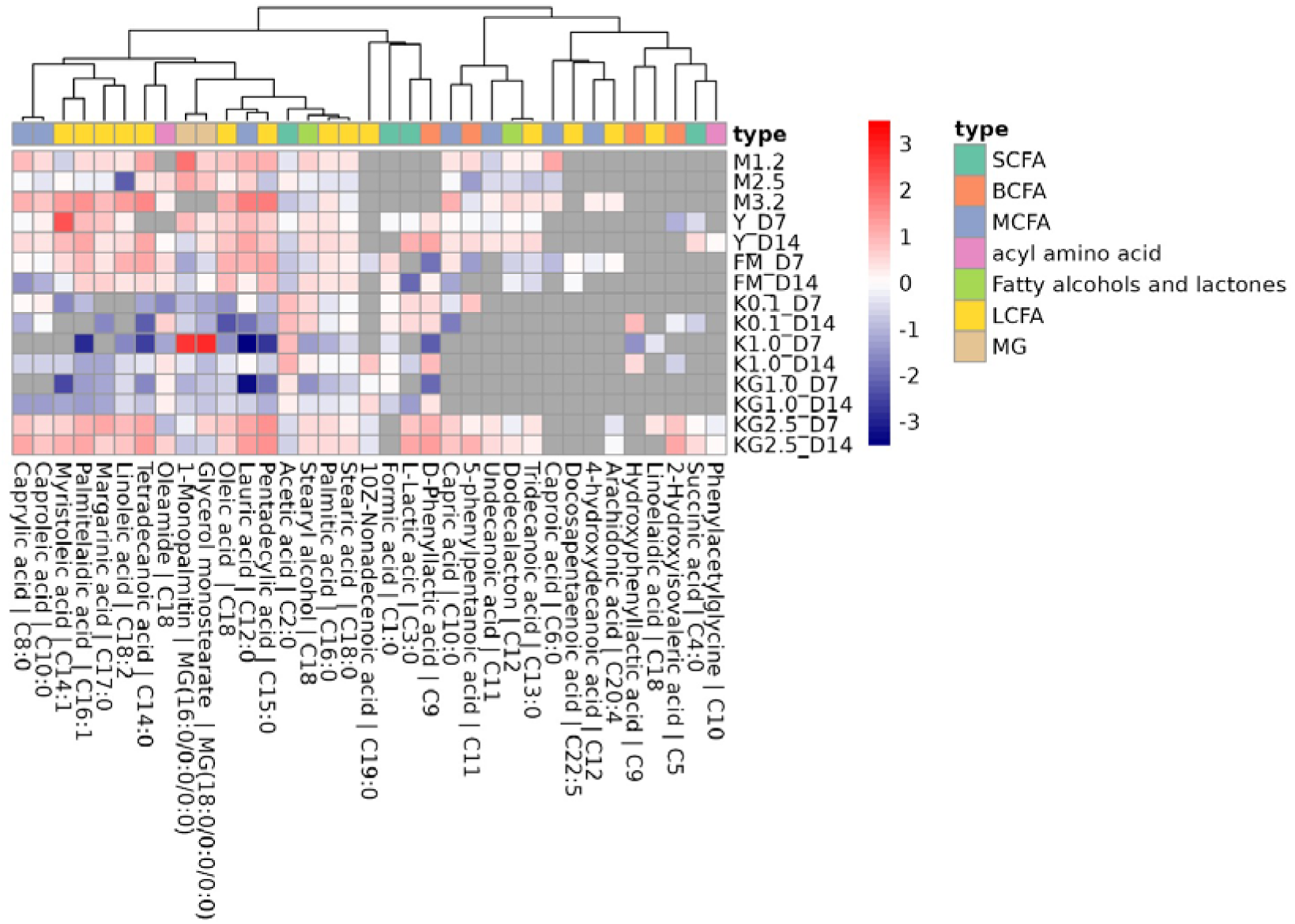
Heatmap of fatty acids concentrations in the products. Succinic acid was produced during fermentation in Y, K and KG but not in FM. Eleven metabolites that were lacked or measured once only are not shown on the graph: eicosapentaenoic acid, nonanoic acid, glutaric acid, dodecyl butyrate, 2-hydroxyisocaproic acid, butanoic acid, propionic acid, malonic acid, nonalactone, 5Z-dodecenoic acid and dihomo-gamma-linolenic acid.

When comparing fermented products with 2.5% fat content with products with 2.5% milk, it was observed that the quantities of medium- and long-chain fatty acids in products with 2.5% fat content were higher than those in milk with the same fat content (Fig. S5).

Notably, the levels of monoglycerides and triglycerides in almost all the products were lower than those found in milk. This may serve as evidence of lipolysis during fermentation. A distinguishing characteristic of fermented foods was the presence of short-chain fatty acids, which were found at higher concentrations than in milk in all fermented milk products.

For product K with a single yeast with 0.1% fat and FM without yeast with 2.5% fat, there was a noticeable decrease in triglyceride content, an increase in monoglycerides and a decrease in free fatty acids. In contrast, for K with 1% fat, Y 1.7%, KG 1%, and KG containing a mix of yeasts with 2.5% fat, there was a decrease in total fat and monoglycerides, accompanied by an increase in long- and medium-chain free fatty acids compared to the rest of the products (see Fig. S6). This observation was likely due to the activities of different sets of yeast and bacterial lipolytic enzymes.

There were more than 500 peaks, after library search and only 27 compounds with majority of fatty acids were identified (see Table S4). Correlation analysis of all identified compounds was performed to assess the possible co-occurrence of the compounds. It can be seen that MCFA and LCFA are clustered into one large cluster, which can be explained by lipolytic processes of decomposition of long chains into shorter ones, as well as the synthesis of protective antimicrobial compounds (Fig. 8). A negative correlation was observed between formic acid and MCFA (e.g. caprylic acid) and LCFA (e.g. undecanoic acid). In contrast, a positive correlation was observed between succinic acid and palmitic acid. Another cluster was formed due to positive correlations between SCFA and BCFA such as L-lactic acid, β-phenyl lactic acid, 2-hydroxy-valeric acid and saturated stearic acid (C18:0). This may indicate that the microbial conversion of these metabolites involves carbohydrates and amino acids. Monoglyceride 1 -monopalmitin was positively correlated with monoglyceride glycerol-stearate, which may be due to specific microbial lipase activity. To interpret the biological significance of such correlations, a more in-depth study of the metabolic pathways of bacteria and yeast present in foods is necessary.

**Figure 8.**
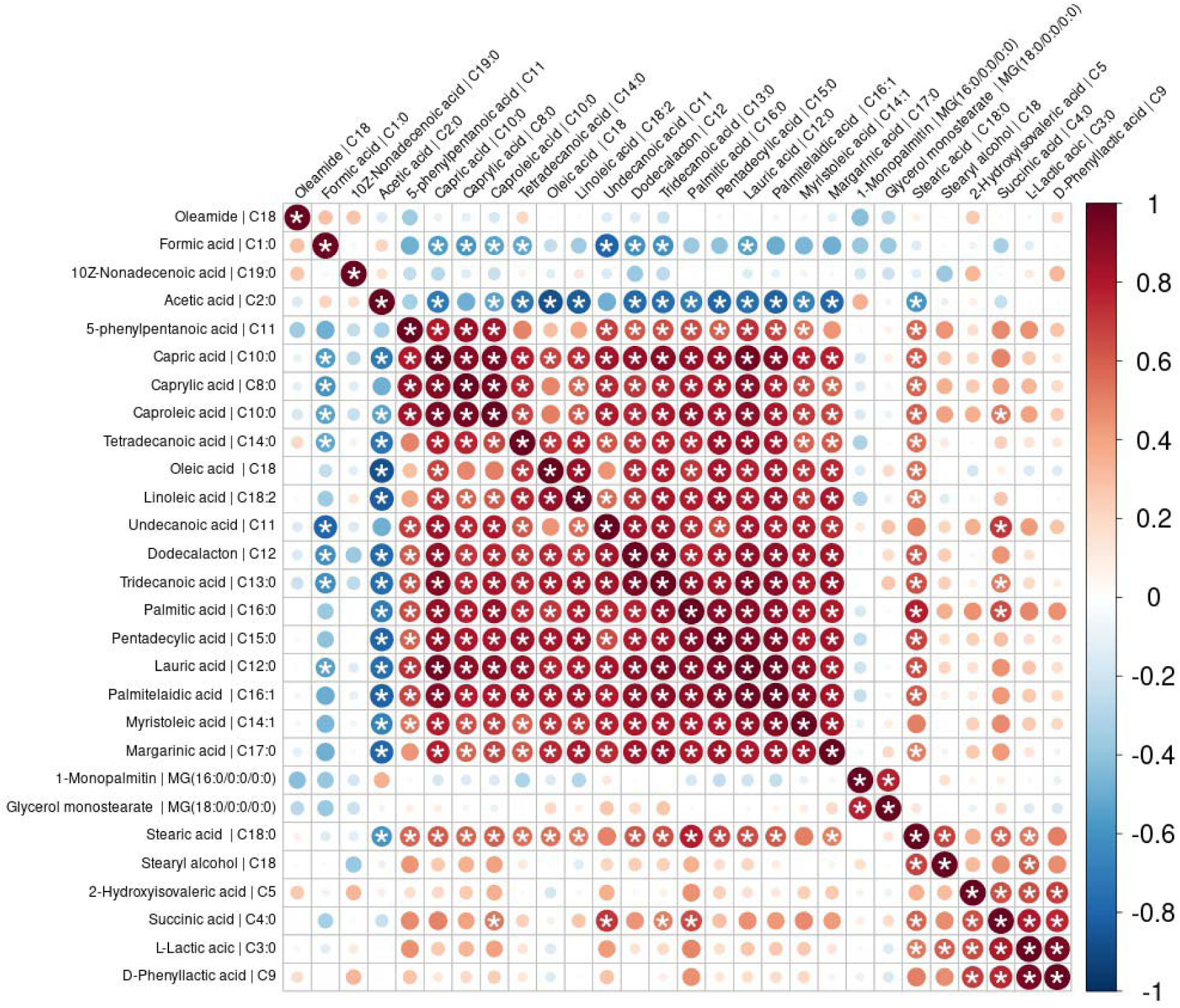
Correlation analysis of fatty acids composition. The values of each metabolite were compared for significance using the Spearman’s correlation coefficient. Negative correlations are shown in blue and positive correlations are shown in red. The diameter of the circles on the heatmap represents larger value correlations. False discovery rate (FDR) corrected *p* values higher than 0.05 are marked with asterisks.

### 3.4. Glycome analysis

The profiles of mono-and disaccharides and polyol myo-inositol were determined using untargeted metabolomics. More than 500 peaks were identified per sample for each product. These peaks were then compared with the library values for simple sugars, resulting in the assessment of values corresponding to 23 standards (see Table S5).

The decomposition products lactose, glucose and galactose were present in the fermented products (Fig. 9). This indicates active lactose decomposition and further metabolism of the monomers during storage. Importantly, there was no accumulation of simple sugars, suggesting that these compounds are actively utilized in other bacterial processes. Overall, the analysis of simple sugar profiles using untargeted metabolomics provides insights into the decomposition and utilization of lactose in fermented milk products. Furthermore, during the life processes of microorganisms, other sugars such as fructose and ribose may also be produced. In addition to sugars, the presence of myo-inositol was identified. Apart from comparing the composition with milk, changes in the levels of simple sugars during storage were evaluated.

**Figure 9.**
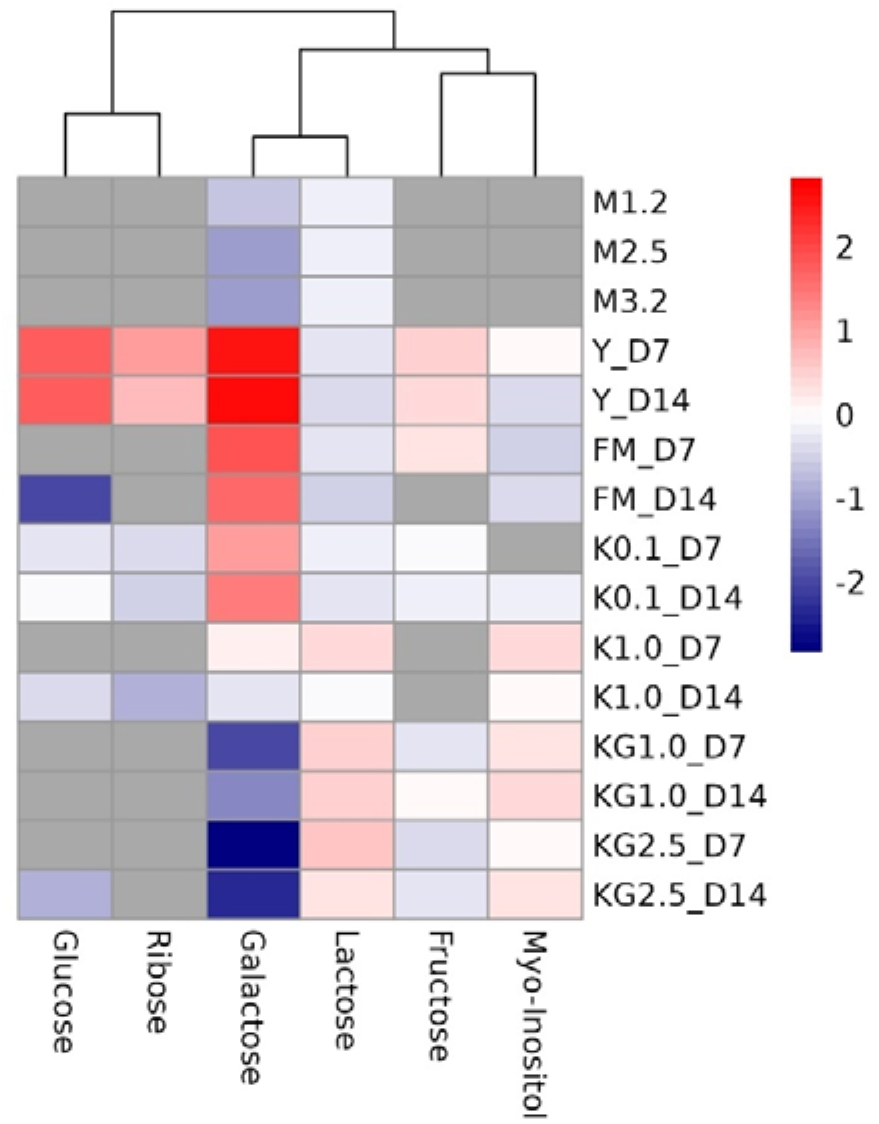
Heatmap shows changes in profile of mono- and disaccharides in fermented milk products during storage (7 vs. 14 days).

There were noticeably higher lactose levels in K and KG than in Y and FM. At the same time, the increase in the amount of different simple sugars occurred differently for all products, probably because of the composition and metabolic potential of the fermenting microbial community. In contrast, an increase in galactose content was observed in Y, FM and K with 0.1% fat, indicating active lactose fermentation by the microbial community during storage.

We observed an accumulation of myo-inositol especially at D14 in all fermented products. This polyol has many protective functions for health and was not detected in our milk samples.

## 4. Discussion

This study was based on untargeted peptidome and metabolome profiling of four types of fermented dairy products and revealed 348 varieties of peptides, 27 fatty acid compounds, and accumulation of BCAAs (valine, leucin, isoleucine), orotic acid and myo-inositol after 14 days of cold storage, resulting from metabolic activity during milk fermentation. The number of peptides and fatty acids decreased slightly, but not significantly, on day 14, in contrast to that on day 7 of storage. We found that 1% of total peptides identified was unique for KG compared to Y, FM and K made with commercial starter cultures. Such specific peptides might be the product of the diverse symbiotic microbial community of wild kefir grains of KG, which includes more than 10 species of yeast, compared to K with a single yeast present (see Table S8). One of the dominant yeast species found in KG was *Malassezia* spp. which accounted for 26% of the total fungal abundance. This yeast might be responsible for the specific metabolite profile, since they are not usually found in kefirs ^41^, which are well characterized ^42^, but are reported to be the primary yeast responsible for the fermentation of kombucha from Brazil ^41^.

The majority of peptides (94%) originated from casein degradation. This was not surprising because the primary source of these peptides is casein, which constitutes up to 80% of the milk protein fraction ^43^. The casein family consists of four main types: alpha S1-casein, alpha S2-casein, beta-casein, and kappa-casein. These caseins are widely present in various food products, ranging from being the primary constituent of cheese to serving as food additives. Although all four casein proteins were related, multiple alignments of their amino acid sequences using the CLUSTAL-Omega program ^44^ indicated minimal homology between the sequences. Each of the four proteins has a distinct amino acid sequence that is unrelated to other casein proteins.

Peptides from other proteins were present in smaller quantities, whereas peptides derived from whey proteins were not detected in this particular study. We did not detect peptides originating from whey proteins α-LA and β-LG, which account for 70-80% of the total whey fraction. Similar results have been reported for other starter cultures, which can be explained by poor proteolysis during fermentation ^45^. In contrast, later studies on kefir from goat milk identified four α-LA peptides and 14 β-LG peptides after 36 h of storage, and none of the α-LA and two β-LG peptides were found in raw goat milk ^46^. In the current study, we analyzed fermented cow dairy products after seven days of storage and did not find any lactalbumin peptides. Their lack might be because: i) few peptides might degrade during prolonged days of storage; ii) the lactalbumin fraction was initially present in cow milk in very small numbers compared to goat-milk; iii) lactalbumins are more resistant to microbial peptidase present in our commercial products, which was also reported previously ^47^ and therefore might be missed by the LC/MS protocol focused on the detection of relatively short peptides.

Based on our study, it appears that the majority of peptides resulting from the proteolysis of beta-casein in fermented milk products were not found in the original milk. Additionally, peptides present in milk were observed at minimal concentrations. This suggests that proteolysis during the fermentation of milk products leads to the generation of unique peptides that are not commonly found in milk. These findings emphasize the significant impact of microbial proteolysis on the composition and characteristics of fermented milk products. In general, kefir had the largest pool of peptides compared to yogurt and fermented milk analyzed in this study, which is in agreement with previous reports ^48^.

Some peptides possess specific physiological functions for human health. In this study, 41 functional peptides were identified. Depending on the peptide composition, they have antihypertensive, antioxidant, bacteriostatic, opioid-like, anti-inflammatory, antiproliferative, antithrombotic, hypolipidemic, hypocholesterolemic, and metal-scavenging properties ^49^. Common examples of very short peptides absorbed by the intestine that have known physiological activity in the blood are valin-proline-proline (VPP) and isoleucine-proline-proline (IPP). Such peptides are formed from milk protein as a result of the protease activity of *Lactobacillus helveticus* and other LAB. These peptides are thermostable, and their resistance to gut proteases helps ameliorate hypertension by inhibiting angiotensin enzymes in the aorta. This effect has been demonstrated in rats ^50^, and later in humans ^49^. Peptides and free amino acids are taken up by enterocytes of the small intestine, and short peptides consisting of 2-3 amino acids are absorbed ^51,52^. However, we were unable to measure such very short functional peptides despite the presence of dipeptide peaks below the resolution levels (see Table S3). Some long peptides with size of 10-51 amino acids (e.g., gonadotropin-releasing hormone 1 and insulin) remain intact, can be absorbed in the intestine, and may have physiological effects on cells. Notably, the physiological effects of enteric peptides are related to their length, and shorter peptides have less effect ^53^. All identified functional peptides in this study belong to the long peptides.

Of note, during fermentation at D14, we found the accumulation of branched chain amino acids (valine, leucine, iso-leucine), phenylalanine, tyrosine, proline, aspartate, pyroglutamic and orotic acid (vitamin B13), whereas tryptophan, glycine, beta-alanine, serine and glutamate were consumed on D7 of product. Thus, the proteolytic activity of the microbial community may lead to the accumulation of certain amino acids from casein ^54^, and BCAA from Greek-style yogurt may elicit different postprandial aminoacidemic responses ^55^. Accumulation of orotate in KG and FM, with the highest concentrations in Y, was observed on D7 and D14, and our results support a single study ^56^, suggesting that microbial synthesis of vitamin B13 might occur under refrigerated conditions despite an initial loss reported for milk under simulated fermentation ^57^.

The composition of the fat fraction, which consists of triglycerides and free fatty acids, was highly dynamic. Longer compounds, such as tri- and monoglycerides, were broken down to free fatty acids. Fatty acids may be broken down into shorter fatty acids; however, the medium-length fatty acid fraction appears to be utilized by microorganisms in their life processes as there was no dramatic accumulation. The short fatty acid fraction was the most highly represented in all fermented products with succinic acid being the most abundant in KG. Lactobacilli present at high levels in dairy products are known for their ability to produce lactate and acetate, and the removal of the bacterial community from the reconstituted kefir community may lead to the persistence of other secondary metabolites ^58^, including succinic acid, which is abundant in carbohydrates ^59^. Kefir consumption was shown to alleviate autism-like behavior in a mouse model of ASD and increase anti-inflammatory Treg cells in the lymph, with increasing succinic acid, elevated *Lachnospiraceae* bacterium A2 taxa, and decreased *Clostridiaceae* abundance in the gut ^60^. Recently, the *Lachnospiraceae* A2 taxon was found to drive IgA levels in the intestine ^61^. Thus, there is strong evidence that kefir metabolites, such as succinic acid, may indirectly contribute to immune protection against pathogenic bacteria in the human gut. Strains of *L. kefiranofaciens,* which are also found in kefir grains of KG but are present in low abundance (0.1%) in the final product, involve upregulation of Treg cells ^62^, consequently inhibiting the secretion of proinflammatory markers but also inducing obesity in high-fat diet (HFD)-fed mice ^63^.

Some functional metabolites including propionic, maleic, butyric, dihomo-gamma linoleic (DGLA) and eicosapentaenoic acid (EPA) were not observed which may indicate their very low concentrations in the analyzed samples. We also did not detect a gut barrier-protective conjugated linoleic acid (CLA), since very low concentrations of this microbial fatty acid as 0.06-0.13 mg/g in the fermented products ^64,65^. The introduction of specific starter cultures, such as *Lactococcus lactis* ssp. *cremoris* MRS 47, may increase CLA concentrations ^66^.

As for carbohydrate components, the fermentation process breaks down lactose into glucose and galactose, which is likely to have a favorable effect on the digestibility of products, especially for people with lactase deficiency, which, as we demonstrated earlier, tends to persist in up to 43% of East Slavs genotypes ^67^. Furthermore, simple sugars such as glucose and fructose were released during lactose fermentation, primarily in Y. These simple sugars may contribute to the overall taste of a product. Overall, glycolysis during fermentation leads to the breakdown of lactose into glucose and galactose, which aids digestibility. It also results in the release of simple sugars and other compounds, contributing to the taste and potential health benefits of fermented products.

Additionally, other sugar-like compounds, such as myo-inositol were released during fermentation. Myo-inositol is component of phytate which is common in cereals and legumes but also can be also released as intermediate product of glucose metabolism by microorganisms ^68^ including primary KG member *K. marxianus* ^69^. To our knowledge, myo-inositol as an end-product has been reported in yogurt from sheep milk ^70^, originating from a raw source. In contrast to that study, we did not observe the source of myo-inositol in cow milk but could see accumulation of this compound in K on day 7 and in K on day 14 with fats >1%, while Y and FM had very low relative quantities. Myo-inositol has a wide range of biological activities, including participation in protection from type 2 diabetes via the reducing blood glucose concentration in metabolic disorders related with insulin resistance, regulation of central nervous system activity, and can potentially serve as pre-biotic for butyrate producing gut microbes ^71,72^. Currently, myo-inositol is being tested as a treatment for disorders of the nervous and reproductive systems and diabetes, including gestational and malformation ^73^ and this polyol was highlighted by meta-review that it can reduce gestational diabetes mellitus ^74^.

In addition to the main components of the products, we found a number of low-molecular weight compounds that are considered to play a role in body functionality. D-phenyl lactic acid, a product of phenylalanine metabolism by bacteria, has antimicrobial activity. Thus, it can be used as an eco-friendly agent for food preservation ^75^. It is also described as a bacterial metabolite capable of specifically modulating the immunity and energy system of the human body through activation of the HCA3 receptor ^76^. The acylated amino acid, oleamide may be formed as a result of the enzymatic degradation of phospholipids. It is an intermediate metabolite that participates in arachidonic acid synthesis catalysed by fatty acid amide hydrolases. This compound has been actively studied as an endocannabinoid ligand to improve sleep quality and psychological health ^77^. Orotic acid is involved in the biosynthesis of pyrimidines and proteins and is essential for the regulation of genes that play a role in the development of cells, tissues, and organisms. It also plays an antioxidant role and is associated with cardiac health (vasorelaxation and cardioprotection) ^78^. The metabolic activity of biotics is the main factor affecting the health impact of consumed fermented products. Among all metabolite classes analyzed in the current study, bioactive peptides remain promising for further detailed studies.

There are several limitations that are important to note. First, the control milk was not the same milk used to produce fermented products. Furthermore, the technological chain of a particular industrial fermented dairy product always involves a mixed formulation consisting of locally normalized milk and skim milk powder, with the aim of adjusting key parameters such as protein and fat for controlled fermentation and nutrition values printed on the label. The composition of metabolites in milk can also vary depending on various factors, such as the conditions under which cows are maintained. This difference in milk used as a reference may introduce some variability in the results obtained during the experiment. Second, non-target or semi-quantitative proteomics methods were used to detect peptides in the sample, but only very rough conclusions could be drawn regarding the concentrations of the detected peptides, indicating whether one peptide is present at higher or lower concentrations than the other. To accurately estimate the absolute quantities of the compounds of interest, it is necessary to perform targeted analysis of sample product batches. Some of the 348 detected peptides were observed at very low concentrations. The number of peptides present in very low concentrations in the samples was beyond the sensitivity of the method.

## Author Contributions

Study conception and design OS and OV; writing – original draft preparation EK; writing – original draft preparation, interpretation of results, review and editing MS; visualization - VO; analysis and interpretation of results - SK; funding acquisition - OV. All authors have read and approved the final manuscript.

## Abbreviations

ACN, acetonitrile; CAA, 2-chloroacetamide; CLA, conjugated linoleic acid; DCNa, sodium deoxycholate; FM, fermented milk; GC-MS, gas chromatography–mass spectrometry; IPP, isoleucine-proline-proline; K, kefir drink; KG, kefir made with wild kefir grains; LAB, lactic acid bacteria; LCFA, long-chain fatty acid; M, normalized milk; MCFA, medium-chain fatty acid ; MSTFA, N-Methyl-N-(trimethylsilyl)-trifluoroacetamide; TCEP, Tris(2-carboxyethyl) phosphine ; TFA, trifluoroacetic acid; UPLC-MS/MS, Ultra-performance liquid chromatography-mass spectrometry; VPP, valine-proline-proline; α-LA, alpha-lactalbumin; β-LG, beta-lactoglobulin;

## Supporting information

full list of peptides

functional list of peptides

amino acids and ornitine

fatty acids

sugars and myo-inositol

total peptides number

fat percentage

microbial compositio of KG

## Acknowledgments

We express our gratitude to the staff of the Lopukhin Federal Research and Clinical Center of Physical-Chemical Medicine for conducting peptidome analysis. We also thank the ABT laboratories, which helped to analyze the profiles of fatty acids, free amino acids and glycome. In addition, we thank the reviewers for their helpful comments and suggestions for improving our manuscript.

## Supplementary Materials

The following supporting information can be downloaded: Supplementary Files 1 and 2, Tables S1-8. Data described in the manuscript are available without restriction at 10.5281/zenodo.14751156 and 10.5281/zenodo.15720835

## TOC graphic

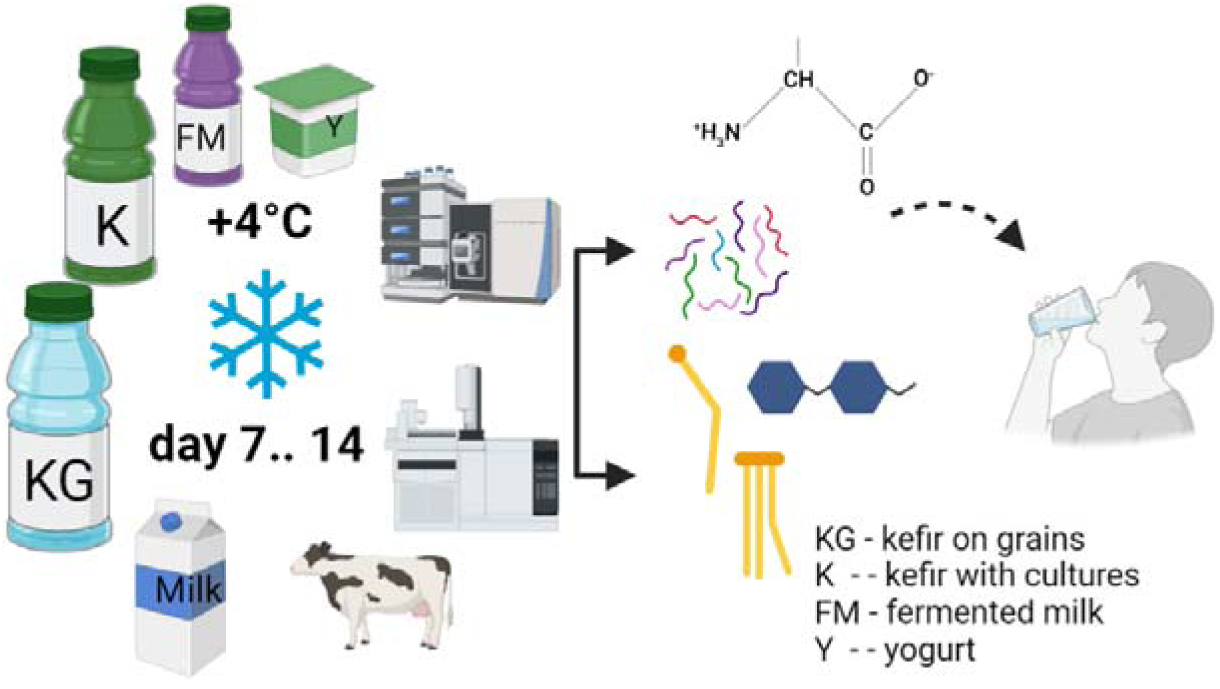

