## Supplementary material for "Peptidome and Metabolome profiling of the fermented milk products revealed accumulation of bioactive compounds during two weeks of cold storage": Supp_1

### Supplementary materials 1

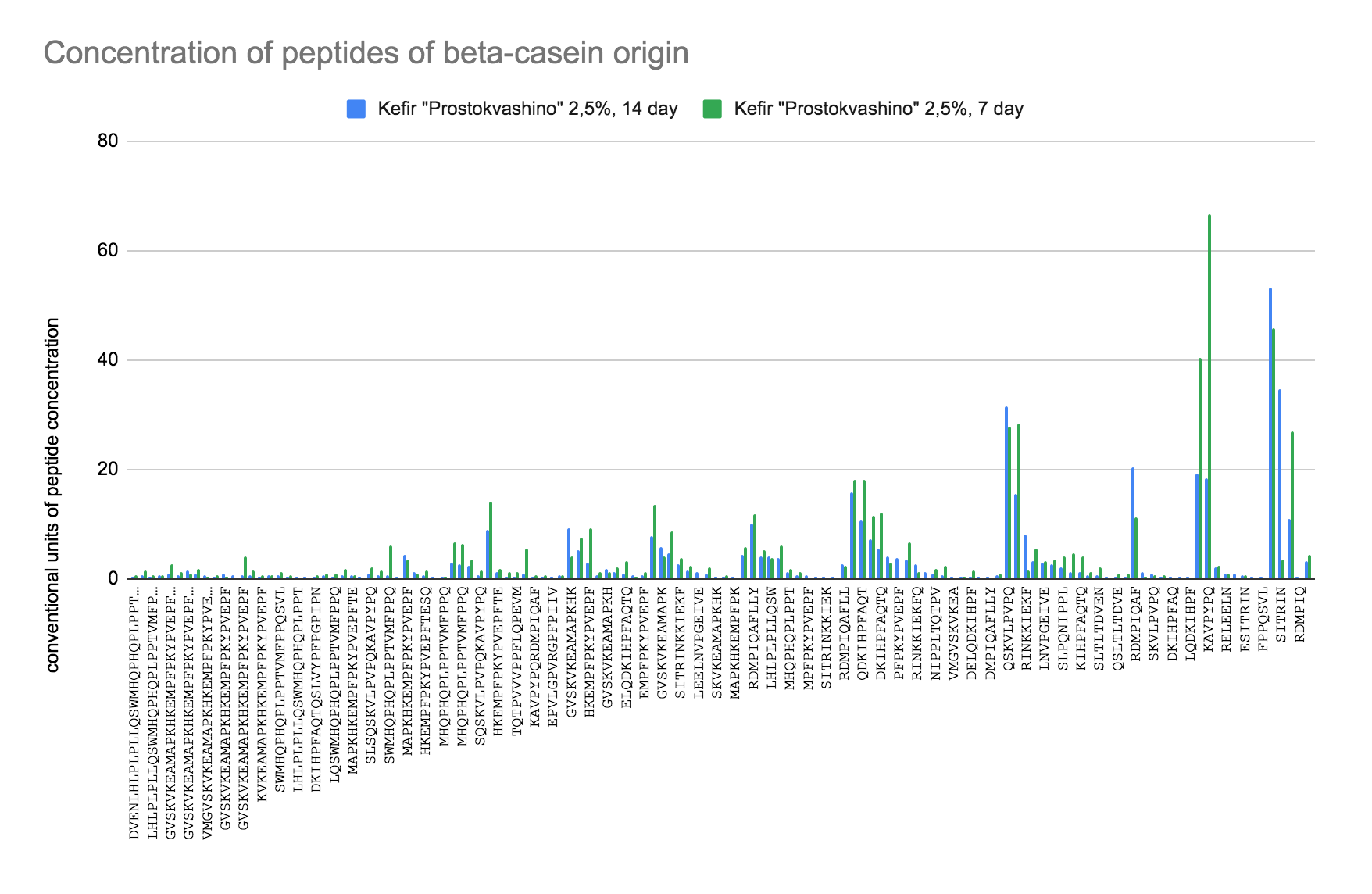

**Figure S1. Concentration of peptides of beta-casein origin. KG2.5_D7 – green colour bars; KG2.5_D14 - blue colour bars.**

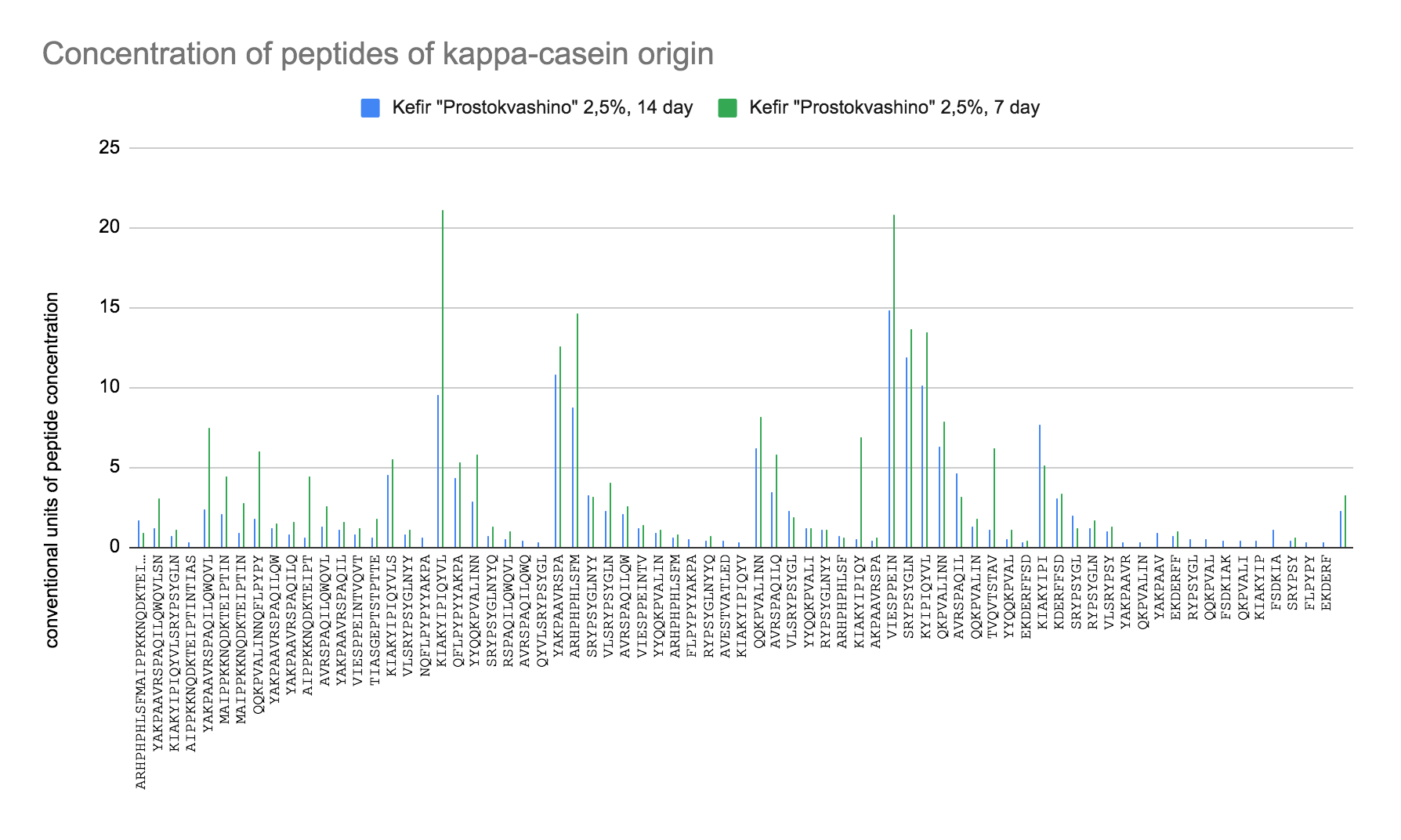

**Figure S2. Concentration of peptides of kappa-casein origin. KG2.5_D7 – green colour bars; KG2.5_D14 - blue colour bars.**

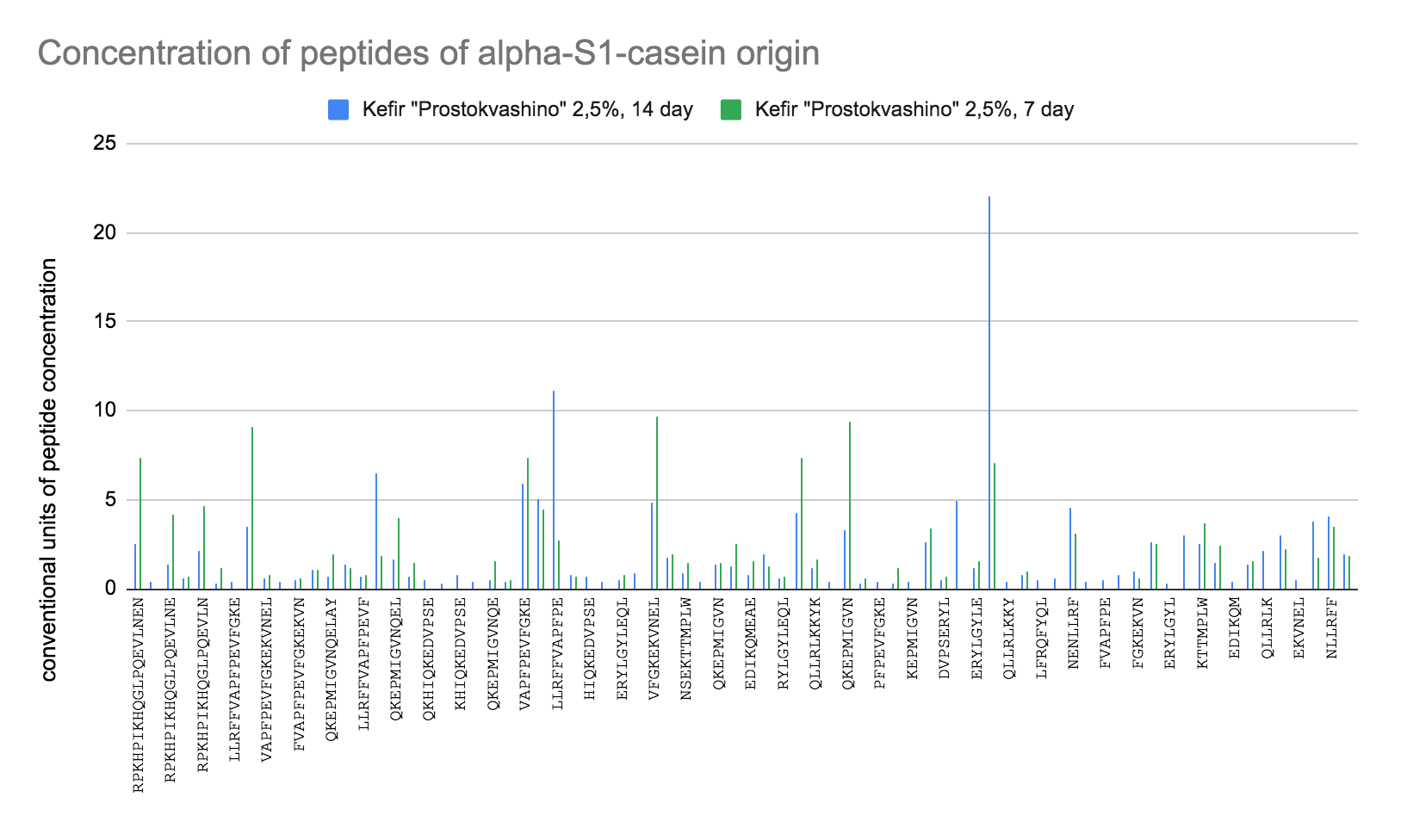

**Figure S3. Concentration of peptides of alpha-S1-casein origin. KG2.5_D7 – green colour bars; KG2.5_D14 - blue colour bars.**

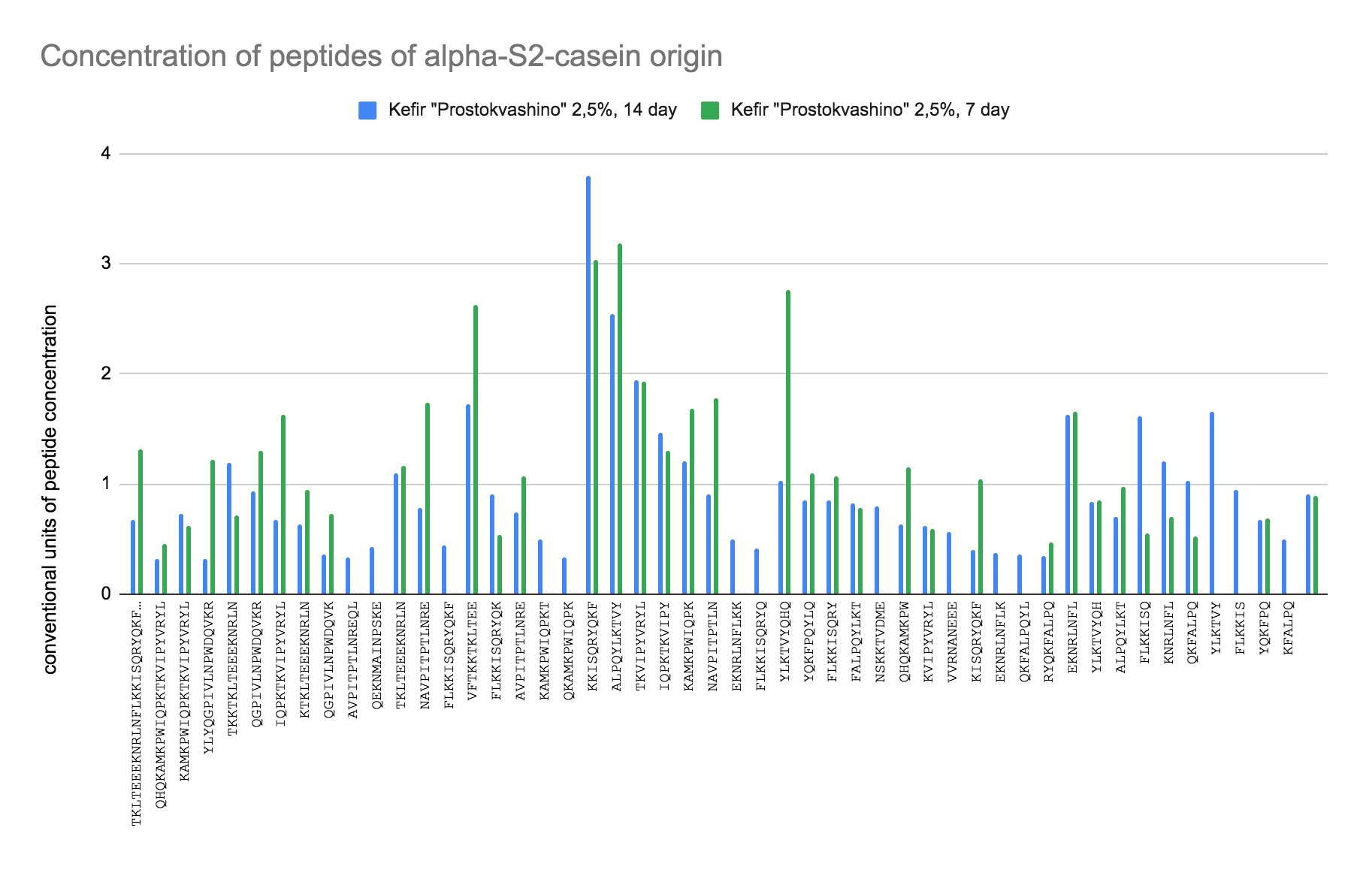

**Figure S4. Concentration of peptides of alpha-S2-casein origin, KG2.5_D7 – green colour bars; KG2.5_D14 - blue colour bars.**

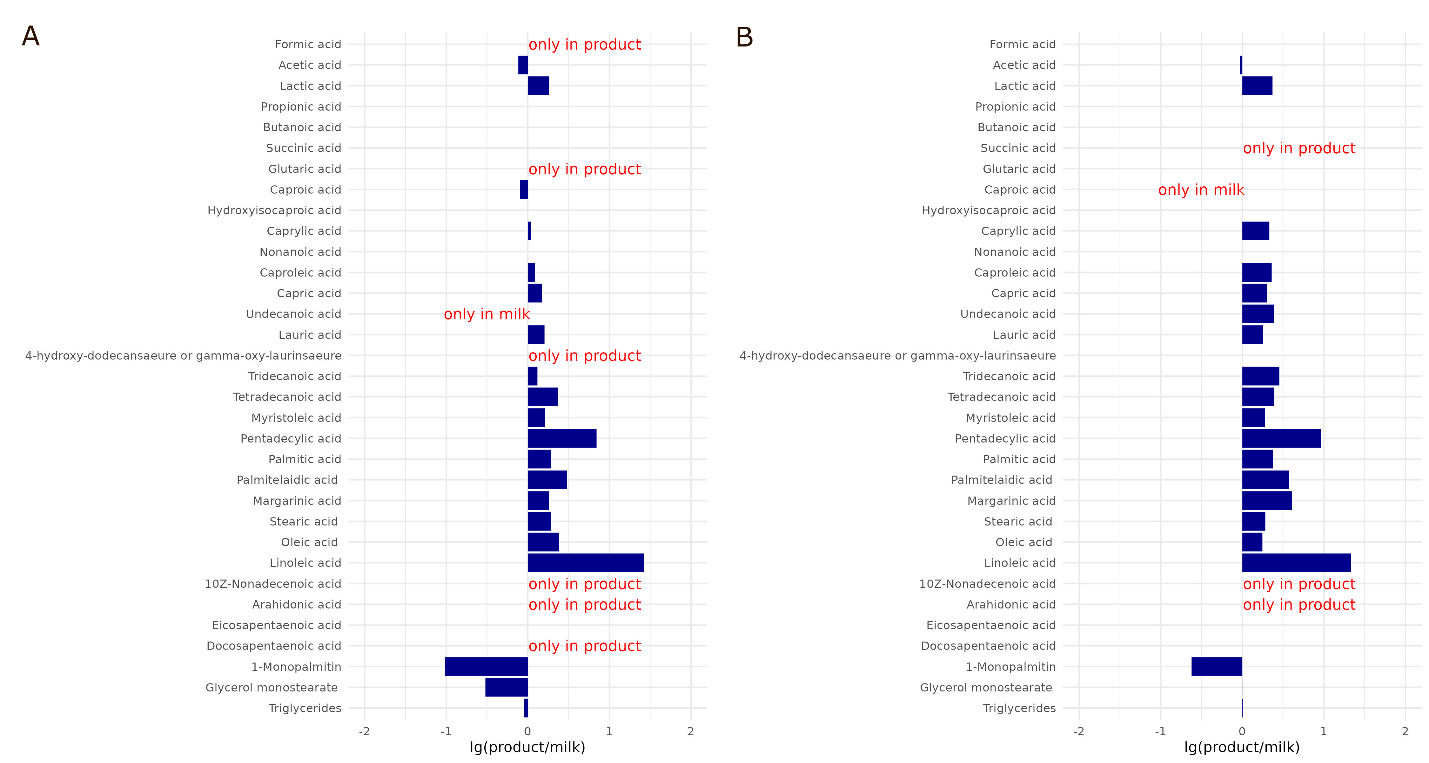

**Figure S5. Comparison of medium and long chain fatty acids content in 2.5% fat products with 2.5% milk (M2.5): (A) FM_D7; (B) KG2.5_D7**

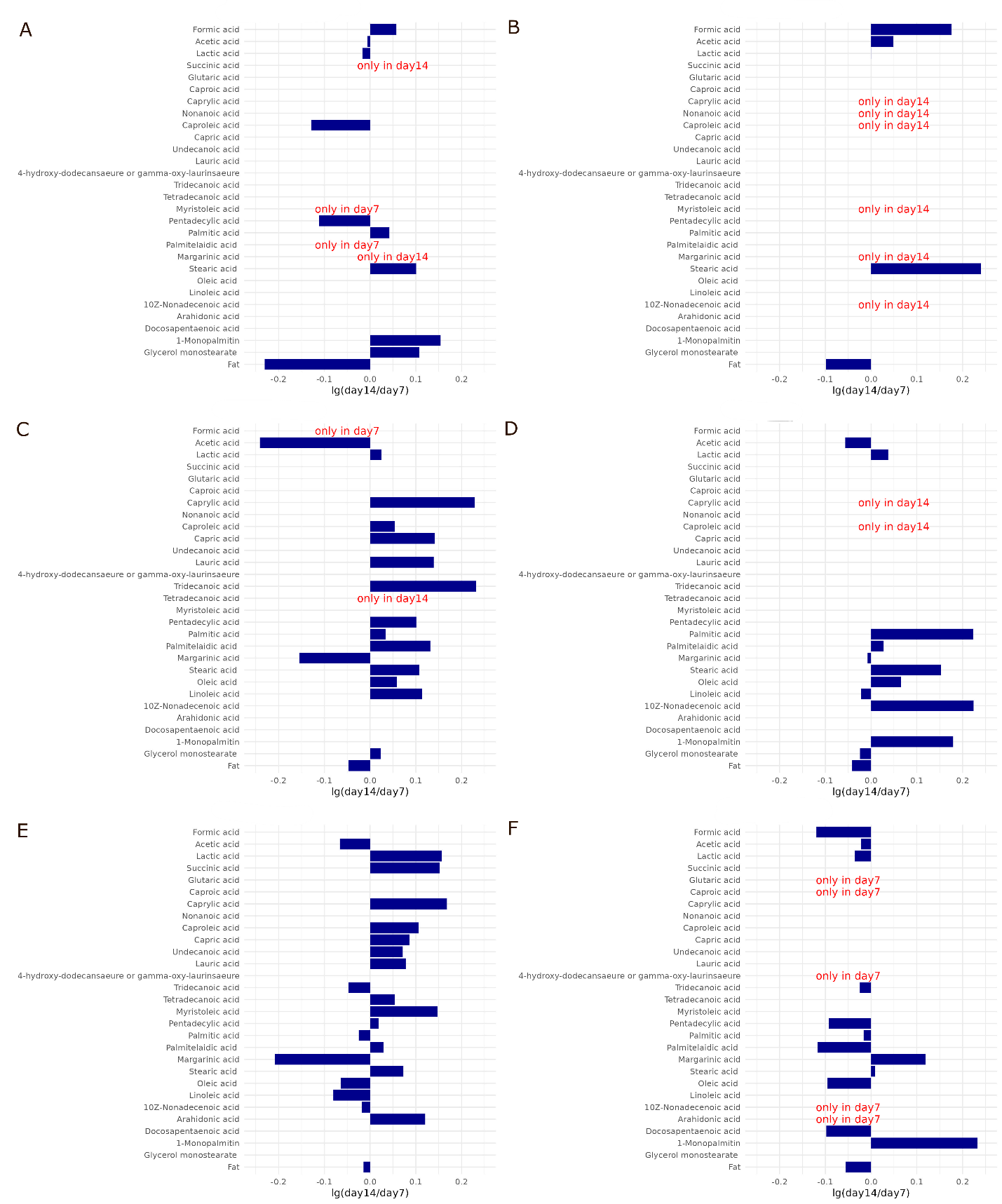

**Figure S6. Comparison of medium and long chain fatty acids content in studied products in 7 and 14 days of storage for (A) K0.1; (B) K1.0; (C) Y; (D) KG1.0; (E) KG2.5; (F) FM**
