## Supplementary material for "Peptidome and Metabolome profiling of the fermented milk products revealed accumulation of bioactive compounds during two weeks of cold storage": Supp_2

**Supplementary Materials 2**

**QC protocols for peptidome and metabolomics**

1. **Calibration standards for peptides**

For each sequence, chemical synthesis of both the target KVLPVPQ and the modified peptide KVLPVPQA was performed by adding an additional amino acid as a non-isotopic tag. The peptides were chromatographically purified and their concentrations were spectrophotometrically measured by absorbance at 214 nm. The molar extinction coefficients of the synthetic peptides were determined using ProtPi (https://www.protpi.ch/Calculator/PeptideTool) with their corresponding amino acid sequences. Final concentrations of purified peptides KVLPVPQ and KVLPVPQA were 0.4 µg/µL and 0.7 µg/µL correspondently.

All synthetic peptides were mixed in a single solution. To construct the calibration curve, a calibration panel of 12 concentrations in 3.3-fold increments was prepared using resulting calibrator by serial dilution. Mass spectrometry analysis was performed on each of the 12 calibration panel samples. Raw data from the analysis of the control samples in *.raw format are available at: 10.5281/zenodo.14751156

For all synthetic peptides, plots of the dependence of peptide MS peak intensities on the actual peptide concentration in the sample were constructed. For all peptide pairs, linear dependence of the two parameters was demonstrated within the range of concentrations used (R-value ≥ 0.95) (see Figure S7).

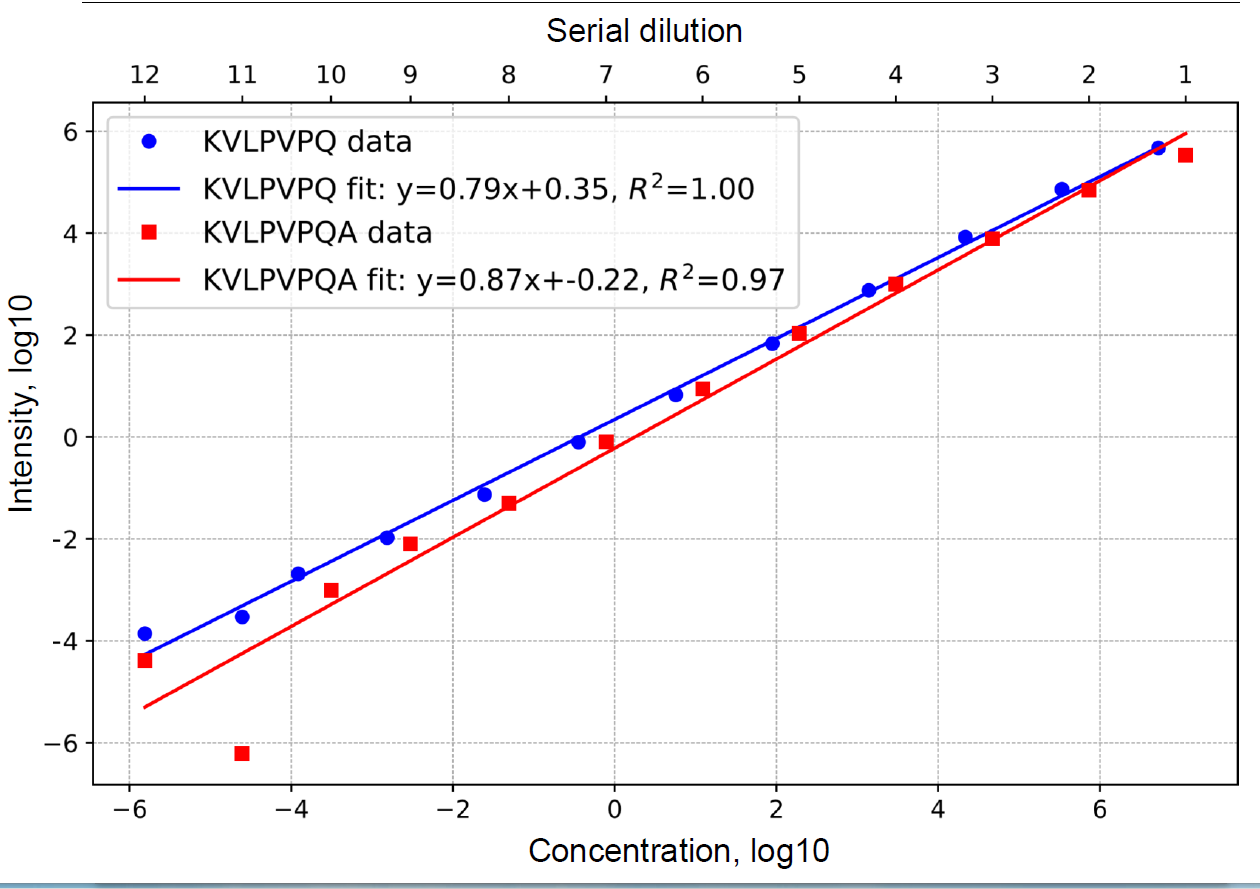

Figure S7. Calibration curve showing intensity vs concentrations of synthetic KVLPVPQ and KVLPVPQ(A) peptides standards

2. **Metabolomic standards and consistency of free fatty acids measurements on different dates.**

Each metabolite peak area was normalized to the internal standard of 2-iso-propylmalic acid (Sigma-Aldrich, cat #333115) for carbohydrates and amino acids; Supelco37 standard (Sigma-Aldrich, cat #CRM47885) for free fatty acids followed by generalized log transformation and data scaling by autoscaling (mean-centered and divided by standard deviation of each variable) as described before ^27^.

Comparative analysis of the relative concentrations of FFAs in milk and fermented milk products. The chromatograms for each sample category are presented below.

In milk samples with 1.5%, 2.5%, and 3.2% fat percentages taken at different dates, there were no changes in the relative concentrations of free fatty acids (see Figures S8-S10).

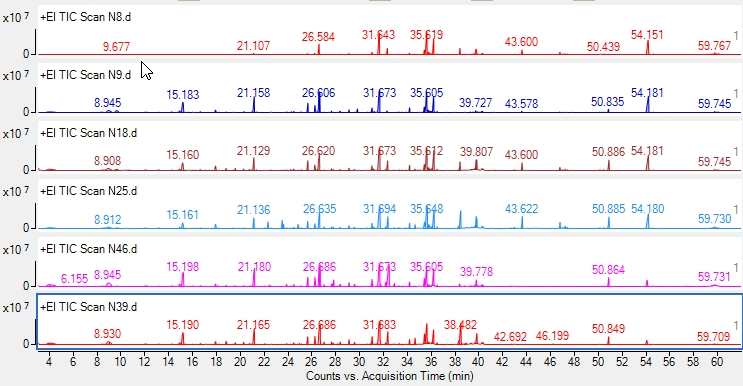

М1.5(4) 11.09.22

М1.5(3) 08.09.22

М1.5(3) 03.09.22

М1.5(1) 01.09.22

М1.5(1) 01.09.22

М1.5(2) 02.09.22

Figure S8. Milk samples with 1.5% fat (M1.5) chromatograms.

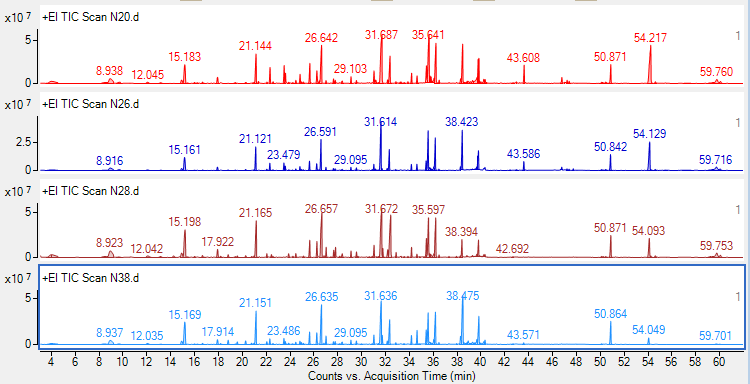

М2.5(4) 11.09.22

М2.5(3) 08.09.22

М2.5(3) 03.09.22

М2.5(2) 02.09.22

Figure S9. Milk samples with 2.5% fat (M2.5) chromatograms.

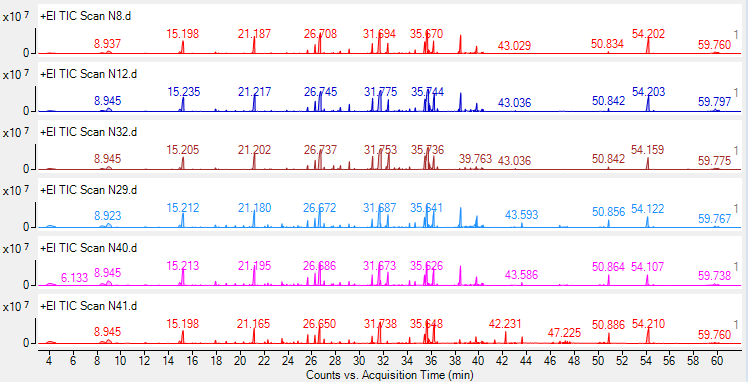

М3.2(4) 11.09.22

М3.2(4) 11.09.22

М3.2(3) 08.09.22

М3.2(3) 03.09.22

М3.2(2) 02.09.22

М3.2(1) 01.09.22

Figure S10. Milk samples with 3.2 % fat (M3.2) chromatograms.

In Kefir with commercial yeast monoculture (K0.1 and K1.0), there were higher amounts of stearic acid and amide of oleic acid on day 14 (D14) than on day 7 (D7) of storage (see Figure S11-S12).

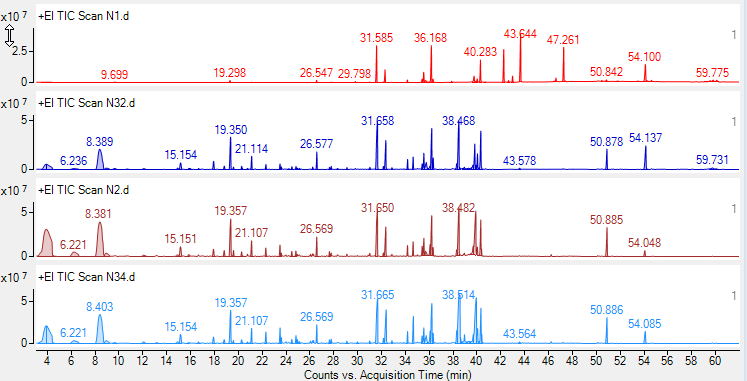

K1.0_D14(2) 08.09.22

K1.0_D7(2) 01.09.22

K1.0_D14(1) 08.09.22

Figure S10. K1.0_D7 and K1.0_D14 chromatograms.

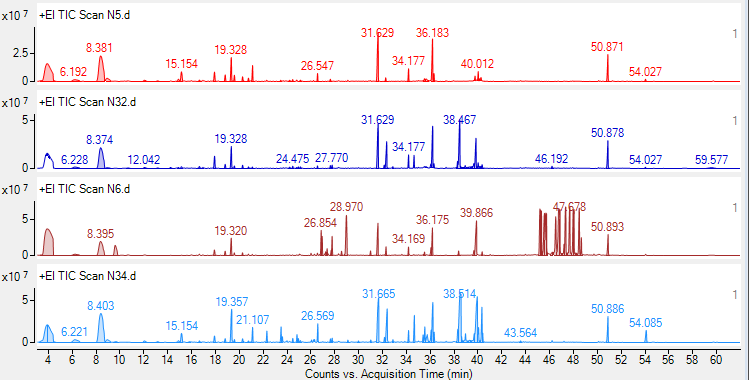

K0.1_D14(2) 08.09.22

K0.1_D14(2) 08.09.22

K0.1_D7(1) 01.09.22

K0.1_D14(1) 08.09.22

Figure S11. K0.1_D7 and K0.1_D14 chromatograms.

For the fermented milk (FM) samples, a slight increase in stearic acid content was observed (see Figure S12).

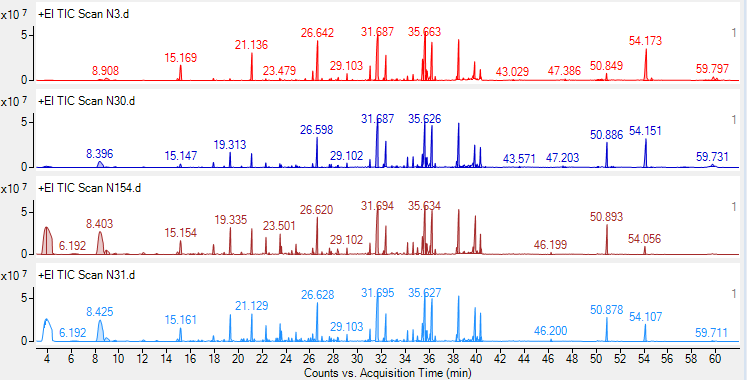

FM_D14(2) 08.09.22

FM_D7(2) 01.09.22

FM_D14(1) 08.09.22

FM_D7(1) 01.09.22

Figure S12. FM_D7 and FM_D14 chromatograms

For kefir on grains with 1% fat (KG1.0), the appearance of new peaks corresponding to arachidonic, decanoic, and 5-phenyl valeric acids with slight decreases in oleic, stearic, and amide oleic acids were observed after 14 days of storage ( Figure S13). In KG2.5, lactic acid, stearic acid, and oleic acid slightly increased during storage, and new peaks corresponding to arachidonic acid and 5-phenyl-valeric acids were observed (see Figure S14).

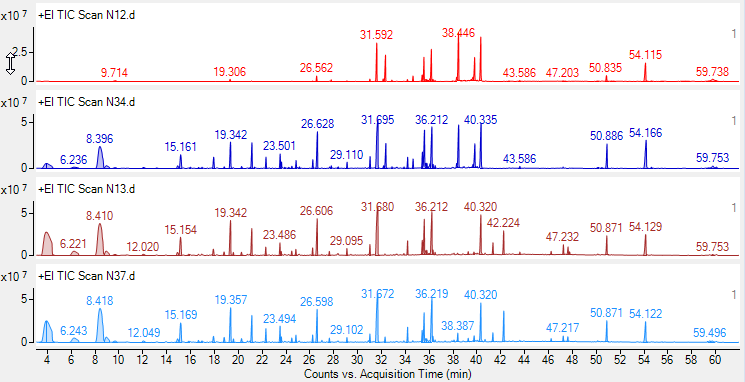

КG1.0_D14(2) 10.09.22

КG1.0_D7(1) 02.09.22

КG1.0_D14(1) 10.09.22

КG1.0_D7(2) 02.09.22

Figure S13. KG1.0_D7 and KG1.0_D14 chromatograms

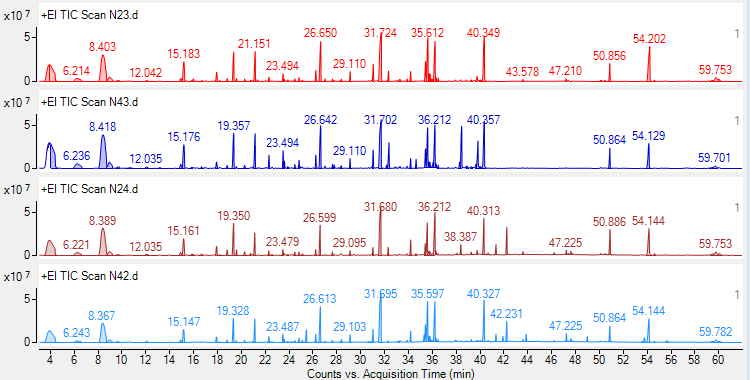

КG2.5_D14(1) 11.09.22

КG2.5_D7(2) 03.09.22

КG2.5_D14(2) 11.09.22

KG2.5_D7 (1) 03.09.22

Figure S14. KG2.5_D7 and KG2.5_D14 chromatograms.

For yogurt samples (Y), there was a profound increase in the relative amounts of lactic, oleic, succinic, and stearic acids, and a slight increase in 5-phenyl valeric acid and glycerol monostearic acid at D14 compared to D7 (see Figure S15).

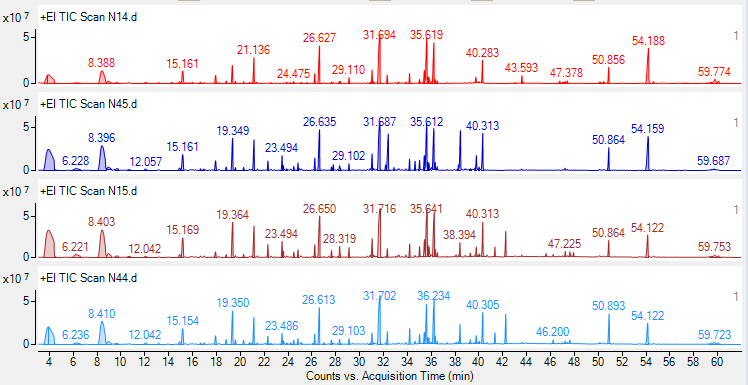

Y_D7(2) 02.09.22

Y_D14(1) 11.09.22

Y_D7(1) 02.09.22

Y_D14 (2) 11.09.22

Figure S15. Y_D7 and Y_D14 chromatograms.
